# Disrupting CD38-driven T cell dysfunction restores sensitivity to cancer immunotherapy

**DOI:** 10.1101/2024.02.12.579184

**Authors:** Or-Yam Revach, Angelina M. Cicerchia, Ofir Shorer, Boryana Petrova, Seth Anderson, Joshua Park, Lee Chen, Arnav Mehta, Samuel J. Wright, Niamh McNamee, Aya Tal-Mason, Giulia Cattaneo, Payal Tiwari, Hongyan Xie, Johanna M. Sweere, Li-Chun Cheng, Natalia Sigal, Elizabeth Enrico, Marisa Miljkovic, Shane A. Evans, Ngan Nguyen, Mark E. Whidden, Ramji Srinivasan, Matthew H. Spitzer, Yi Sun, Tatyana Sharova, Aleigha R. Lawless, William A. Michaud, Martin Q. Rasmussen, Jacy Fang, Claire A. Palin, Feng Chen, Xinhui Wang, Cristina R. Ferrone, Donald P. Lawrence, Ryan J. Sullivan, David Liu, Uma M. Sachdeva, Debattama R. Sen, Keith T. Flaherty, Robert T. Manguso, Lloyd Bod, Manolis Kellis, Genevieve M. Boland, Keren Yizhak, Jiekun Yang, Naama Kanarek, Moshe Sade-Feldman, Nir Hacohen, Russell W. Jenkins

## Abstract

A central problem in cancer immunotherapy with immune checkpoint blockade (ICB) is the development of resistance, which affects 50% of patients with metastatic melanoma^1,2^. T cell exhaustion, resulting from chronic antigen exposure in the tumour microenvironment, is a major driver of ICB resistance^3^. Here, we show that CD38, an ecto-enzyme involved in nicotinamide adenine dinucleotide (NAD^+^) catabolism, is highly expressed in exhausted CD8^+^ T cells in melanoma and is associated with ICB resistance. Tumour-derived CD38^hi^CD8^+^ T cells are dysfunctional, characterised by impaired proliferative capacity, effector function, and dysregulated mitochondrial bioenergetics. Genetic and pharmacological blockade of CD38 in murine and patient-derived organotypic tumour models (MDOTS/PDOTS) enhanced tumour immunity and overcame ICB resistance. Mechanistically, disrupting CD38 activity in T cells restored cellular NAD^+^ pools, improved mitochondrial function, increased proliferation, augmented effector function, and restored ICB sensitivity. Taken together, these data demonstrate a role for the CD38-NAD^+^ axis in promoting T cell exhaustion and ICB resistance, and establish the efficacy of CD38 directed therapeutic strategies to overcome ICB resistance using clinically relevant, patient-derived 3D tumour models.

---

Cancer immunotherapy with immune checkpoint blockade (ICB) has revolutionised cancer treatment, providing durable responses and cures for a subset of patients with advanced metastatic disease^4^. Despite the success of ICB in melanoma and other cancers, ICB resistance remains a central challenge and is ultimately fatal^5^. Currently, no approved therapies exist for patients with innate or acquired resistance to ICB. Recently, a phase II clinical trial evaluating CTLA-4 +/- PD-1 blockade in anti-PD-1 refractory melanoma patients demonstrated modest objective response rates (ORR) for ipilimumab (9% ORR) monotherapy versus ipilimumab plus nivolumab (28% ORR) with high rates of grade 3 or higher immune-related adverse effects (35% and 57%, respectively)^6^, underscoring the need for renewed focus on the development of rational therapeutic strategies to overcome ICB resistance.

Tumour-infiltrating CD8^+^ T lymphocytes (TILs) are key determinants of anti-tumour immunity and successful response to ICB^7–9^. Single-cell sequencing has offered deeper insights into intratumoural T cell heterogeneity, revealing that CD8^+^ T cells defined by expression of stem-like transcription factors (e.g., *TCF7*) are enriched in melanoma patients sensitive to ICB, whereas exhausted CD8^+^ T cells, defined by high expression of co-inhibitory receptors, are enriched in tumours from ICB-resistant melanoma patients^10^. Exhausted T cells develop during chronic antigen stimulation and are characterised by loss of proliferative potential and diminished effector function, and may acquire immunosuppressive properties^9–12^. Therapeutic strategies directed at promoting stem-like T cell expansion, enhancing effector functions, and preventing (or reversing) T cell exhaustion would represent rational approaches to overcome ICB resistance.

CD38, an ecto-enzyme involved in NAD^+^ catabolism, is upregulated during T cell activation and, more recently, has been associated with terminal T cell exhaustion in cancer and chronic viral infection^13,14^. CD38^hi^CD8^+^ T cells are induced following unsuccessful response to ICB contributing to ICB resistance, and depletion of CD38^hi^CD8^+^ cells improved anti-tumour immunity in murine tumour models^15^. While several studies have shown that genetic deletion or pharmacologic inhibition of CD38 improves anti-tumour immunity^16,17^, the precise role of the CD38-NAD^+^ axis in governing the balance of tumour infiltrating CD8^+^ T cell states is not known. Importantly, the potential impact of targeting CD38 to overcome ICB resistance in human tumour models has not been tested.

Here we show that upregulation of CD38 in exhausted CD8^+^ cells is strongly associated with ICB resistance, and that targeting CD38 overcomes resistance to ICB in both murine and patient-derived tumour models. We show that CD38^hi^CD8^+^ T cells exhibit features of T cell exhaustion, with low *TCF7* expression, decreased proliferative capacity, diminished effector function and altered bioenergetics. Further, we show that pharmacologic inhibition or genetic deletion of CD38 can reverse these metabolic defects promoting *TCF7* expression, improved T cell function, and augmentation or restoration of ICB sensitivity.

## CD38^+^CD8^+^ TILs predict ICB resistance

We first examined the expression of CD38 in tumour-infiltrating CD8^+^ T cells using published single-cell RNA sequencing (scRNAseq) data of immune cells from patients with advanced melanoma treated with ICB^10^ (**Fig. 1a, Extended Data Fig. 1a-b**). We observed increased *CD38* expression in CD8^+^ T cell clusters associated with T cell exhaustion (clusters 1-3, **Fig. 1a-b**), which closely mirrored the expression of PD-1 (*PDCD1*) (**Fig. 1b-c**). CD8^+^ T cell states associated with early T cell activation or effector memory function (clusters 4-6) were marked with low *CD38* expression and high expression of the transcription factor *TCF7* (**Fig. 1d**). As *TCF7*-expressing CD8^+^ cells are associated with ICB response^10^, we sought to determine if *CD38*-expressing CD8^+^ T cells were associated with ICB resistance. Consistent with prior observations^10,15^, we observed a higher proportion of tumour-infiltrating *CD38* expressing CD8^+^ T cells in ICB non-responders (ICB-NR) in melanoma (**Fig. 1e-f**). As previously shown^15^, CD8-specific expression of *CD38* also demonstrated high predictive power for ICB resistance in melanoma patients (AUC 0.87, *P* = 1.53 x 10^-5^) (**Fig. 1g**), and higher proportions of *CD38* expressing CD8^+^ T cells were observed in both baseline (pre-treatment) and post-treatment tumours from ICB-resistant melanoma patients, although the observations were more robust in post-treatment specimens (**Extended Data Fig. 1c-d**). Furthermore, CD8^+^ T cell-specific *CD38* expression was associated with resistance both to single-agent PD-1 blockade and dual PD-1/CTLA-4 blockade (**Extended Data Fig. 1e-f**).

**Figure 1.**
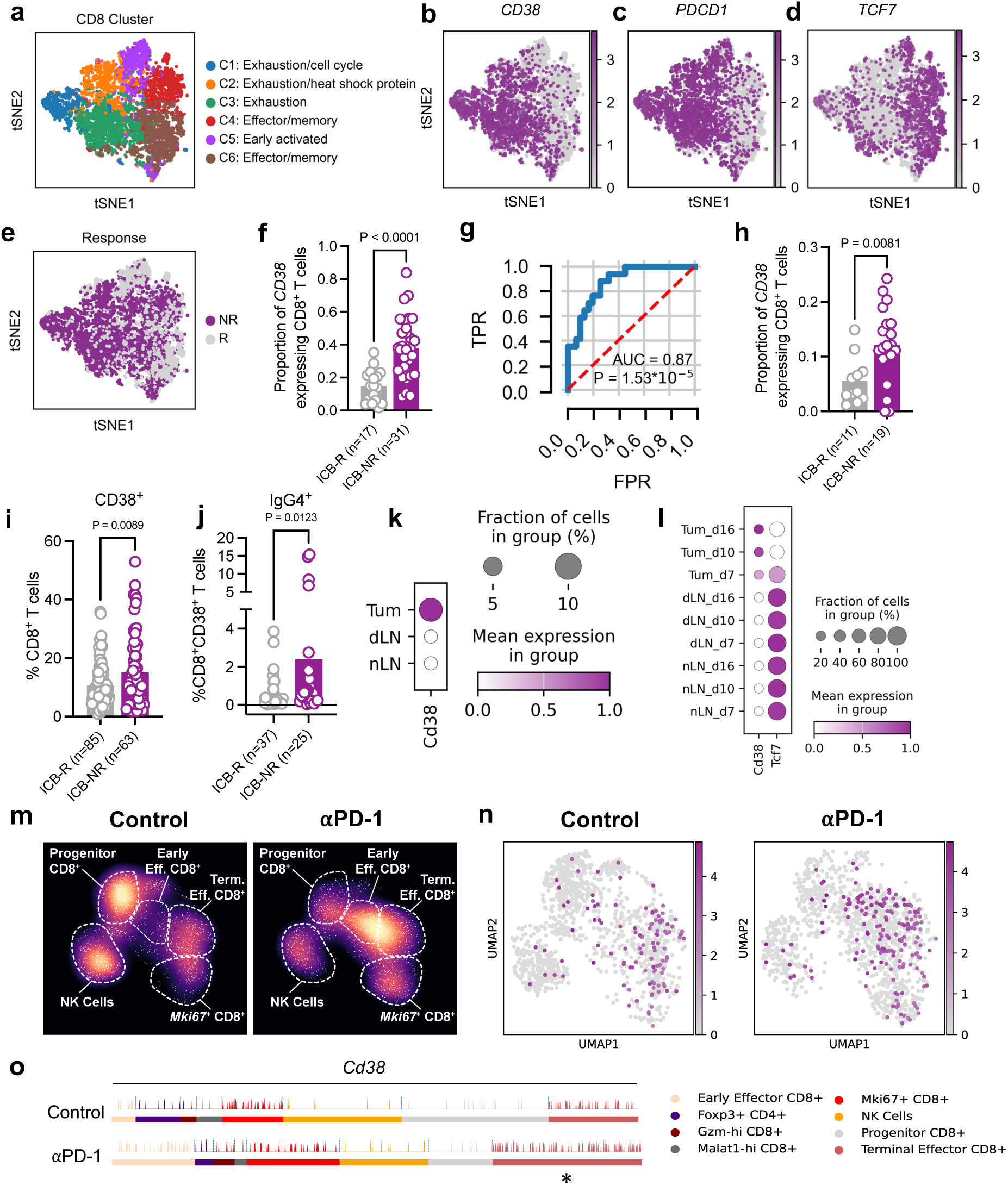
CD38^+^ CD8^+^ T cells are associated with ICB resistance. **a-g**, scRNAseq of CD45^+^ immune cells from melanoma patients^10^ with (**a**) t-SNE plot of CD8^+^ T cell clusters (n=6,350); (**b-d**) t-SNE plot of CD8^+^ T cell clusters showing expression of **(b)** *CD38*, **(c)** *PDCD1*, and **(d)** *TCF7*; **(e)** t-SNE plot of CD8^+^ T cell clusters by ICB response; **(f)** Proportion of *CD38* expressing CD8^+^ T cells from indicated groups. Responders (ICB-R) and non-responders (ICB-NR). Means (bars) and individual values (open circles) are shown. 2-sided unpaired *t*-test; (**g)** Receiver Operating Characteristic (ROC) curve, demonstrating the predictive power of CD38^+^CD8^+^ T cells for ICB resistance. **h**, Validation cohort evaluating proportion of *CD38* expressing CD8^+^ T cells from indicated groups; 2-sided unpaired *t*-test. **i-j**, CyTOF analysis of peripheral blood examining levels of **(i)** CD38^+^CD8^+^ T cells and **(j)** IgG4^+^ CD38^+^CD8^+^ T cells from ICB responding and non-responding patients; Means (bars) and individual values (open circles) are shown; 2-sided unpaired *t*-test. **k-l**, Dot plots indicating (**k**) *Cd38* expression in CD8^+^ TILs from B16-ova tumours (Tum), tumour draining lymph nodes (dLN), and normal lymph nodes (nLN) and (**l**) expression of *Cd38* and *Tcf7* (Tum, dLN, and nLN) at day 7, 10 and 16^20^. **m-o**, scRNAseq of CD3^+^ TILs from IgG control and αPD-1 treated mice bearing B16-ova tumours^21^ showing (**m**) UMAP of T/NK cell clusters by condition; (**n**) *Cd38* gene expression in UMAP from **(m)**; (**o**) tracks plots of *Cd38* gene expression by T/NK cluster. 2-way ANOVA with Sidak correction for multiple comparisons. Statistics in Extended Data Fig. 2i. \**P* < 0.05.

Sub-cluster analysis of immune populations in melanoma tumours demonstrated that the exhausted CD8^+^ T cells (Cluster 6 in Extended Data Fig. 1a) showed the highest predictive ability for ICB resistance (AUC 0.85, *P* = 3.09 x 10^-4^), compared to the exhausted/cell cycle cluster (Cluster 11, Extended data in Fig. 1a, AUC 0.63, *P* = 0.091) and exhausted/heat shock cluster (Cluster 9, Extended data in Fig. 1a, AUC 0.62, *P* = 0.415) (**Extended Data Fig. 1g-i)**. *CD38* expression in memory T cells, cytotoxic T cells, lymphocytes, regulatory T cells (Tregs), B cells, plasma cells, monocytes/macrophages, and dendritic cells also showed lower predictive power for ICB resistance, and were mostly not significant (**Extended Data Fig. 1j)**. While tumour-specific expression of *CD38* has been observed in NSCLC and other cancers^16^, CD38 expression in melanoma tumour cells was rarely observed in CD45-cells (mostly tumour) from melanoma samples, compared to CD45^+^ immune cells (**Extended Data Fig. 1k).**

To validate these findings, an independent cohort of melanoma patients treated with ICB was evaluated by scRNAseq analysis of tumour infiltrating CD8^+^ T cells (**Extended Data Fig. 1l, Supplementary Table 1**). Enrichment of *CD38*-expressing CD8^+^ T cells was again observed in ICB-NR patients (*n*=19) compared to ICB-R patients (*n*=11) (**Fig. 1h)**, and expression of *CD38* and *TCF7* were observed in distinct T cell clusters, with CD38 expression enriched in exhausted T cell clusters (**Extended Data Fig. 1m-n**). To determine if the association between *CD38* expressing CD8^+^ T cells and ICB resistance could be observed in other cancers, we examined the expression of *CD38* in CD8^+^ T cells from patients with non-small cell lung cancer (NSCLC)^18^. Consistent with our observations in melanoma, *CD38* expression in CD8^+^ T cells was associated with lack of ICB treatment benefit in NSCLC and was predictive for lack of response (AUC 0.75, *P* = 0.002) (**Extended Data Fig. 2a-b)**. Taken together, these observations indicate that *CD38*-expressing tumour infiltrating CD8^+^ T cells are strongly associated with ICB resistance in multiple cancer types.

## Enrichment of CD38^+^CD8^+^ T cells during tumour progression

Higher circulating levels of CD38^+^CD4^+^ T cells have been associated with diminished ICB sensitivity^11^. To determine if elevated CD38 expression in CD8^+^ T cells in peripheral blood was also associated with ICB resistance, we performed multiplexed cytometry by time of flight (CyTOF) of CD8^+^ T cells in melanoma patients’ blood in an independent cohort before and after initiating ICB treatment (**Supplementary Table 2**). Levels of peripheral CD38^+^CD8^+^ T cells were significantly higher in melanoma patients with ICB resistance (**Fig. 1i**). Interestingly, pre-treatment circulating levels of CD38^+^CD8^+^ T cells were not significantly different between responders and non-responders, whereas higher levels of CD38^+^CD8^+^ T cells were observed in post-treatment samples of ICB non-responders (**Extended Data Fig. 2c-d**). Increased IgG4^+^ was also observed on peripheral blood CD38^+^CD8^+^ T cells in ICB non-responders, indicating that these cells were bound by the clinically administered IgG4 isotype anti-PD-1 antibodies^19^ which was not observed in baseline samples (**Fig. 1j, Extended Data Fig. 2e**). In agreement with these observations, circulating plasma levels of CD38 were increased in melanoma patients 6 weeks after initiating ICB treatment, but remained elevated at 6 months only in non-responder (ICB-NR) patients **(Extended Data Fig. 2f)**, indicating that CD38^+^CD8^+^ T cells can be detected in the peripheral blood following ICB treatment and are also associated with ICB resistance.

To determine the temporal dynamics of CD8^+^ T cell-specific CD38 upregulation in tumours, we examined time-resolved scRNAseq data from B16 murine melanoma tumours with matched tumour draining lymph nodes (dLN) and normal lymph nodes (nLN)^20^. While *Cd38* expression was observed in B cells in tumours, dLN, and nLN, *Cd38*-expressing CD8^+^ T cells were observed only within tumours (**Fig. 1k, Extended Data Fig. 2g**). Expression of *Tcf7* in CD8^+^ T cells was observed largely in nLN and dLN, and relative expression was decreased in tumour infiltrating CD8 T cells (**Fig. 1l**). Further, while modest residual *Tcf7* expression was observed in smaller tumours at early time points (D7), *Tcf7* was absent by D10 and remained undetectable at D16 as tumour burden increased (**Fig. 1l**). In contrast, *Cd38* expression was largely absent in nLN and dLN, but was detectable in tumours by D7 with increased expression at D10 and D16 with increasing tumour burden (**Fig. 1l)**. Therefore, while CD38^+^CD8^+^ T cells can be detected in the periphery, CD38 upregulation and the parallel loss of *Tcf7* expression in CD8^+^ T cells occurs specifically in the tumour microenvironment during tumour progression.

## ICB treatment increases CD38^+^CD8^+^ T cells

Ineffective ICB treatment promotes induction of dysfunctional CD8^+^ T cells that contribute to ICB resistance in murine tumour models^15^. To further evaluate the effect of ICB on *Cd38* expression in CD8^+^ T cells, we examined scRNAseq data of tumour infiltrating CD45^+^ immune cells from B16-ova tumours^21^. Aggregated data from isotype (IgG) control and anti-PD-1 treated mice was used to perform clustering and to examine the relative abundance of T/NK cell populations (**Extended Data Fig. 2h**). Anti-PD-1 treatment resulted in the expansion of early effector CD8^+^ T cells and terminal exhausted/effector CD8^+^ T cells with a reduction of progenitor CD8^+^ T cell population (**Fig. 1m**). *Cd38* expression was significantly increased only in terminal exhausted/effector CD8^+^ T cells (**Fig. 1n-o, Extended Data Fig. 2i-l**). Thus, while *Cd38* is expressed on various T/NK cell populations (and multiple immune cell types), its upregulation following ICB treatment is observed primarily in exhausted CD8^+^ T cells.

## CD38^hi^ CD8^+^ T cells are dysfunctional

We next sought to examine the association between *CD38* and T cell exhaustion. Increased expression of multiple co-inhibitory receptors (*PDCD1*, *HAVCR2*, *ENTPD1*, *CTLA4*, *LAG3*) was observed in *CD38*-high CD8^+^ T cells from ICB treated melanoma patients, whereas *CD38* expression was comparatively low in CD8^+^ T cells expressing *TCF7* (**Fig. 2a)**^10^. Analysis of coexpression deviation proportions (CDP) in our validation melanoma scRNAseq data set (see **Fig. 1h**) demonstrated co-expression of *ENTPD1* (encoding CD39), *PDCD1* (encoding PD-1), *CTLA4*, *HAVCR2* (encoding TIM-3), and *LAG3* with *CD38* in CD8^+^ TILs, and an inverse correlation with *TCF7* expression (**Fig. 2b)**. Examination of differentially expressed genes in CD38^+^ and CD38^-^ tumour infiltrating CD8^+^ T cells from melanoma patients treated with ICB^10^ revealed increased expression of co-inhibitory receptors (e.g., *PDCD1*, *HAVCR2*) in CD38^+^ CD8 T cells, and increased expression of markers of stem/memory T cells (e.g., *TCF7*, *IL7R*, *CCR7*) in CD38^-^ CD8^+^ T cells (**Fig. 2c, Supplementary Table 3)**. Analysis of tumour-infiltrating T cells from B16-ova tumours^21^ revealed similar upregulation of co-inhibitory receptors and exhaustion-related molecules in CD38^+^ tumour infiltrating T cells, whereas increased expression of *Tcf7*, *Il7r,* and *Ccr7* was observed in CD38^-^ TILs (**Fig. 2d, Supplementary Table 4)**.

**Figure 2.**
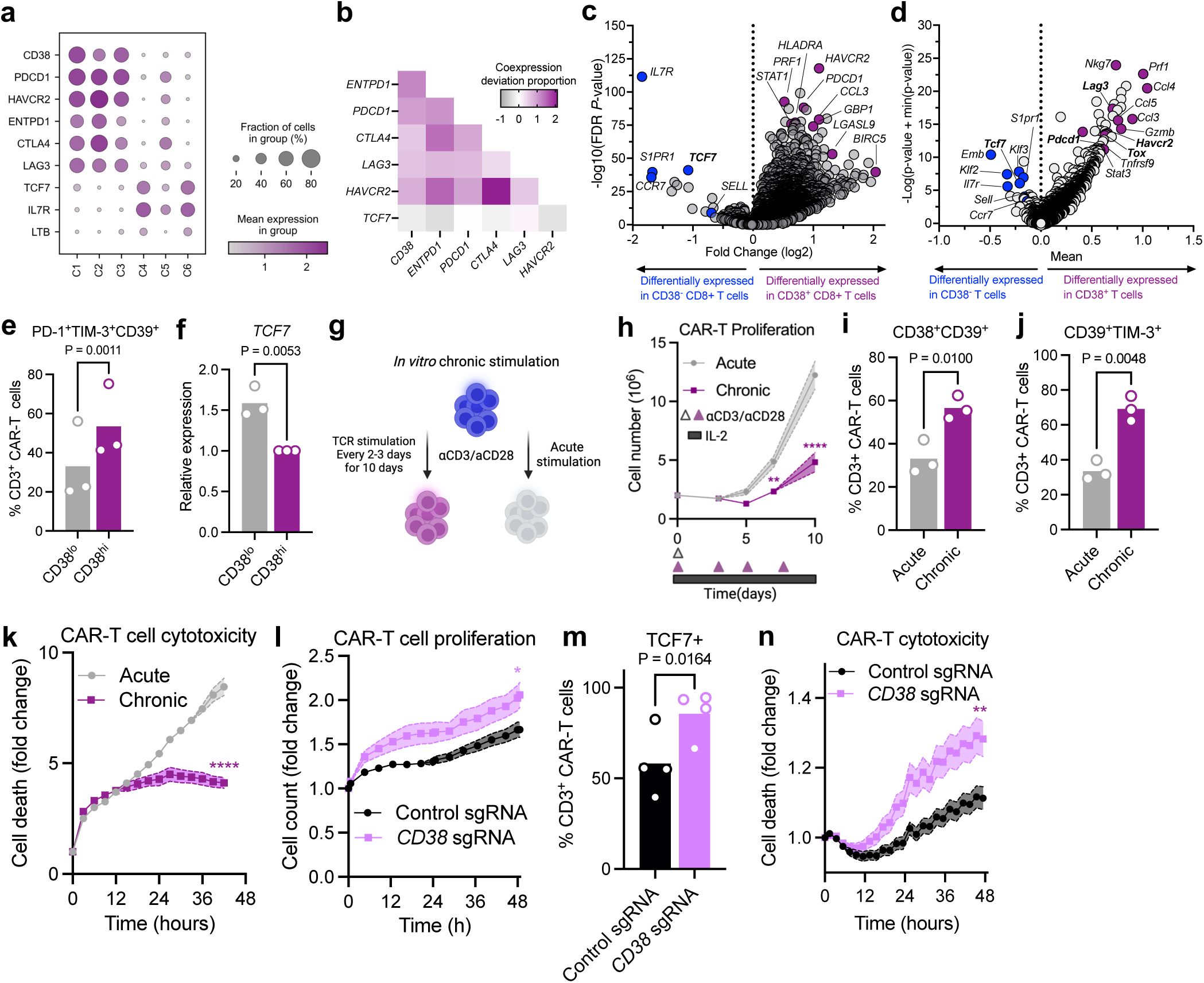
CD38^hi^CD8^+^ T cells are dysfunctional. **a**, Dot plot showing the expression of exhaustion and effector/memory-related genes in CD8^+^ TILs from human melanoma tumours^10^. **b**, Co-expression deviation proportion plot demonstrating co-expression of exhaustion-related genes and *TCF7* from melanoma validation cohort. **c-d**, Volcano plots depicting differentially expressed genes based on CD38 expression in (**c**) CD8^+^ TILs from human melanoma^10^ and (**d**) CD3^+^ TILs from B16-ova murine melanoma^21^. **e-f**, Surface staining (**e**) for PD-1^+^CD39^+^TIM-3^+^ (n=3; 2-sided paired *t*-test) and (**f**) *TCF7* expression (n=3; 2-sided unpaired *t*-test) in sorted CD38^hi^ and CD38^lo^ B7-H3.CAR-T cells. Means (bars) and individual values (open circles) are shown. **g**, Scheme depicting acute and chronic TCR stimulation. **h-k**, Acute and chronic B7-H3.CAR-T (**h**) proliferation assay; Means +/- s.e.m. (shaded areas) are shown (*n*=3; 2-way ANOVA with Sidak correction for multiple comparisons). Surface staining for CD38^+^CD39^+^ **(i)** and CD39^+^TIM-3^+^ (**j**); Means (bars) and individual values (open circles) are shown (*n*=3, 2-sided paired *t*-test) **(k)** cytotoxicity assay towards 10164 patient derived melanoma cell line. Representative experiment out of three is presented. Means +/- s.e.m. (shaded area) are shown (*n*=3 biological replicates; 3 independent experiments; 2-way ANOVA with Sidak correction for multiple comparisons). **l-n**, analysis of chronically stimulated control sgRNA and *CD38* sgRNA B7-H3.CAR-T cells; **(l)** Proliferation assay; Means +/- s.e.m. (shaded area) are shown (*n*=3 biological replicates; 3 independent experiments; 2-way ANOVA with Sidak correction for multiple comparisons); (**m**) TCF7 intracellular staining. Means (bars) and individual values (open circles) are shown (n=4; 2-sided paired *t*-test;); (**n)** Cytotoxicity assay 10164 melanoma cells. Means +/- s.e.m. (shaded area) are shown (n=3 biological replicates; 3 independent experiments; 2-way ANOVA with Sidak correction for multiple comparisons). \**P* < 0.05, \*\**P* < 0.01, \*\*\*\**P* < 0.0001.

To validate these findings, we turned to human B7-H3-directed chimeric antigen receptor T-cells (B7-H3.CAR-T)^22,23^. CD38^hi^ and CD38^lo^ sub-populations of B7-H3.CAR-T cells were isolated by fluorescence-activated cell sorting (FACS), and consistent with our melanoma TIL scRNAseq data, CD38^hi^ CAR-T cells exhibited increased expression of exhaustion related surface markers (PD-1, TIM-3, CD39) and decreased expression of *TCF7* and *FOXO1* compared to CD38^lo^ B7-H3.CAR-T cells (**Fig. 2e-f, Extended Data Fig. 3a-c**). To model the development of T cell dysfunction, we performed repetitive *in vitro* T cell receptor (TCR) stimulation (every 2-3 days) with αCD3/αCD28 beads for 10-14 days (“chronic” stimulation) compared to a one time activation with αCD3/αCD28 beads followed by IL-2 expansion (“acute” stimulation) (**Fig. 2g**). Chronic TCR stimulation of CAR-T cells resulted in reduced proliferative capacity **(Fig. 2h)**, increased levels of CD38^+^CD39^+^ and CD39^+^TIM-3^+^ cells **(Fig. 2i-j, Extended Data Fig. 3d)**, and reduced ability to lyse target cancer cells compared to standard (acute) stimulation (**Fig. 2k, Extended Data Fig. 3e-g),** recapitulating key features of T cell exhaustion. Similar observations were obtained with patient derived CD8^+^ TILs and matched primary tumour cells **(Extended Data Fig. 3h-i**).

## CD38 loss prevents T cell dysfunction

To determine if CD38 is necessary for the development of T cell dysfunction, we deleted *CD38* in B7-H3.CAR-T cells using CRISPR/Cas9 gene editing (**Extended Data Fig. 3j)**. *CD38* sgRNA B7-H3.CAR-T cells displayed a clear proliferative advantage over control sgRNA CAR-T cells (**Fig. 2l)**, accompanied by an increased proportion of TCF7^+^ CAR-T cells (**Fig. 2m**). Interestingly, surface expression of CD39, TIM-3, and PD-1 were largely unaffected by CD38 loss suggesting that despite strong co-expression with these genes, CD38 does not directly regulate the expression of exhaustion markers (**Extended Data Fig. 3k-l**). Further, *CD38* sgRNA CAR-T cells exhibited increased cytotoxic capacity towards patient-derived melanoma cell lines compared to control sgRNA CAR-T cells (**Fig. 2n)**, indicating that *CD38* deletion reduces T cell dysfunction in an *in vitro* model of T cell exhaustion driven by chronic TCR stimulation.

## CD38 blockade overcomes ICB resistance

Given that upregulation of CD38 in CD8^+^ T cells was associated with T cell dysfunction and ICB resistance, and that CD38 deletion was able to restore T cell effector function, we next examined CD38 blockade as a strategy to overcome ICB resistance. To examine the effect of CD38 blockade using a clinically-relevant model of human cancer, we performed *ex vivo* profiling of patient-derived organotypic tumour spheroids (PDOTS)^21,24,25^, a biomimetic technology for studying tumour-immune dynamics using living tumour biopsies grown in 3D microfluidic culture. PDOTS derived from patients with melanoma and select other cancers were treated with anti-PD-1 (pembrolizumab) +/- anti-CD38 (daratumumab)^26^ compared to untreated control (**Fig. 3a**). We observed a significant reduction in PDOTS viability following dual PD-1/CD38 blockade, which demonstrated superior anti-tumour activity compared to anti-PD-1 or anti-CD38 alone in a PDOTS cohort (*n*=30) largely comprised of tumours from ICB-resistant melanoma patients (**Fig. 3b, Extended Data Fig. 4a, Supplementary Table 5**). Examination of patient-specific responses to PD-1 +/- CD38 blockade revealed a 20% response rate (6 of 30) to anti-CD38 treatment and a 56.7% response rate to dual PD-1/CD38 blockade (17 of 30) compared to a 3.3% response rate to single-agent PD-1 blockade (1 of 30) (**Fig. 3c**, **Extended Data Fig. 4a**). Exceptional *ex vivo* response to dual PD-1/CD38 blockade was observed in several PDOTS from patients with ICB-resistant tumours (52% of responders, 9 of 17), including PDOTS 10101 and 10213 derived from patients with ICB-resistant cutaneous and mucosal melanoma, respectively (**Fig. 3d-e, Extended Data Fig. 5a-b**).

**Figure 3.**
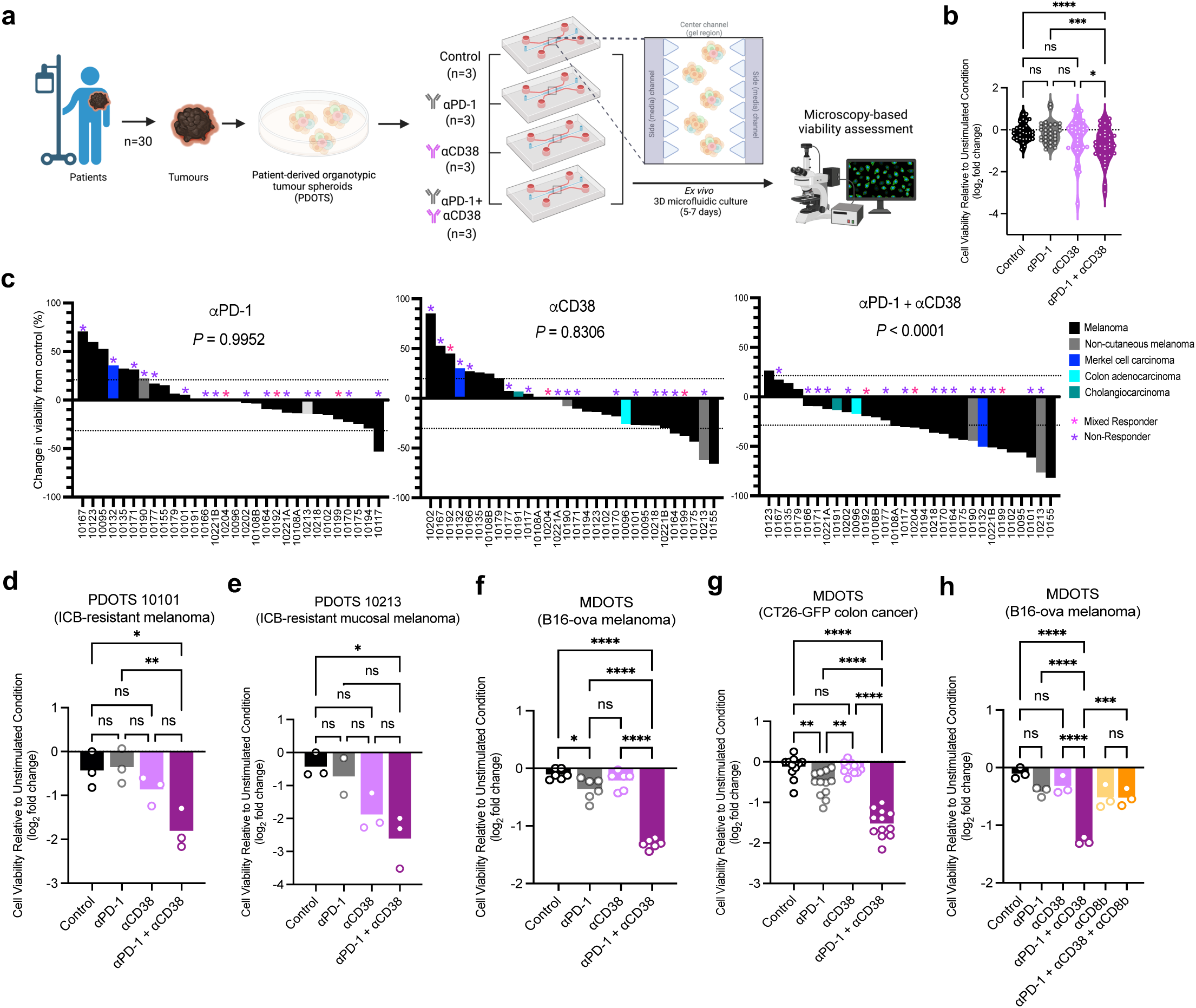
CD38 blockade overcomes ICB resistance. **a**, Scheme of PDOTS preparation. **b**, Violin plot of PDOTS (*n*=30) viability assessment following treatment with anti-PD-1 (250 *μ*g/mL pembrolizumab), anti-CD38 (200 *μ*g/mL), or combined anti-PD-1+anti-CD38. Individual values (open circles) indicate mean for each PDOTS specimen; one-way ANOVA with Tukey correction for multiple comparisons. **c**, waterfall plots for PDOTS (*n*=30, indicated tumour types) with indicated treatments. Response is defined as 30% reduction from control (dashed lines). **d-e**, PDOTS viability assessment from (**d**) ICB-resistant cutaneous melanoma (10101) and (**e**) ICB-resistant mucosal melanoma (10213) with indicated treatments (*n*=3 biological replicates per PDOTS specimen, one-way ANOVA with Tukey correction for multiple comparisons). **f-g**, Viability assessment of (**f**) B16-ova MDOTS (n=6 biological replicates, 2 independent experiments), (**g**) CT26-GFP MDOTS (n=12 biological replicates, 4 independent experiments), (**h**) B16-ova MDOTS (n=3 biological replicates, 1 independent experiment) with indicated treatments. Means (bars) and individual values (open circles) are shown (one-way ANOVA with Tukey correction for multiple comparisons. \**P* < 0.05. \*\**P* < 0.01; \*\*\**P* < 0.001; **** *P* < 0.0001; *ns*, not significant.

Multispectral flow cytometry analysis (prior to PDOTS culture) and immunofluorescence staining of PDOTS in 3D microfluidic culture confirmed expression of CD38 and PD-1 on the surface of tumour infiltrating immune cells (**Extended Data Fig. 5c-d)**. *Post hoc* multivariate analysis of multispectral flow cytometry data and selected clinical data failed to identify any statistically significant association with PDOTS response, although CD8^+^PD-1^+^ and CD8^+^CD38^+^ T cells were most strongly associated with sensitivity to dual PD-1/CD38 blockade (**Extended Data Fig. 6a-c)**.

To determine if CD38 blockade could overcome ICB resistance in murine tumour models, we performed *ex vivo* profiling of murine-derived organotypic tumour spheroids (MDOTS)^24^. Given the accumulation of CD38^+^CD8^+^ T cells with larger tumours (after 10 days; see **Fig. 1l**), MDOTS were prepared using day 10-14 tumour explants to enrich CD38^+^CD8^+^ T cells and CD38^+^PD-1^+^CD8^+^ T cells (**Extended Data Fig. 6d**). B16-ova and CT26-GFP MDOTS exhibited modest response to single-agent PD-1 or CD38 blockade, but showed significantly improved tumour control with combined PD-1/CD38 blockade in both models (**Fig. 3f-g, Extended Data Fig. 6e-f**). Further, we demonstrated that CD8^+^ T cell activity was required for the combinatorial effect of dual PD-1/CD38 blockade in B16-ova MDOTS (**Fig. 3h**). Lastly, *in vitro* CD38 blockade in isolated CD8^+^ TILs from CT26-GFP murine tumours increased the proportion of IL-7Rα^+^CD8^+^ stem-cell-like T cells while decreasing CD69^+^CD8^+^ T cells (**Extended Data Fig. 6g-i)**^8,10,27^ (see Fig. 2d). Taken together, these findings demonstrate that CD38 blockade can overcome ICB resistance in human and murine tumour models.

## CD38^+^ T cells exhibit altered bioenergetics

To identify key cellular processes associated with T cell specific upregulation of CD38, we next performed gene set enrichment analysis (GSEA) using tumour infiltrating CD38^+^CD8^+^ T cells from ICB-treated melanoma patients^10^ and tumour infiltrating CD38^+^ T cells from B16-ova tumours^21^. Top scoring gene sets included oxidative phosphorylation and interferon α/γ (IFNα/γ) response, which correlated strongly between mouse and human data (**Fig. 4a, Extended Data Fig. 7a-d, Supplementary Tables 6-7**). Given these findings and previous reports linking CD38 to altered mitochondrial fitness^14,28^, we examined the mitochondrial mass and mitochondrial membrane potential (MMP) of CD38^lo^ and CD38^hi^ T cells. Increased mitochondrial mass and MMP were observed in CD38^hi^ B7-H3.CAR-T cells, human melanoma CD8^+^ TILs, and Jurkat T cell leukaemia cells (**Extended Data Fig. 7e-k**). Moreover, deletion of *CD38* resulted in reduced mitochondrial mass and lower MMP compared to control sgRNA following chronic TCR stimulation of B7-H3.CAR-T cells (**Extended Data Fig. 7l-m**). *CD38* sgRNA CAR-T cells also exhibited increased rates of oxygen consumption (OCR) and extracellular acidification (ECAR) (**Fig. 4b, Extended Data Fig. 7n**), indicating improvements in mitochondrial respiration and glycolysis, respectively. OCR and maximal respiratory capacity were also increased in CAR-T cells treated with CD38i (**Fig. 4c**), mirroring the effect of genetic deletion of CD38. Increased maximal respiratory capacity (following addition of the mitochondrial uncoupling agent, FCCP) suggests improved T cell fitness under conditions of increased metabolic stress in *CD38* sgRNA CAR-T cells.

**Figure 4.**
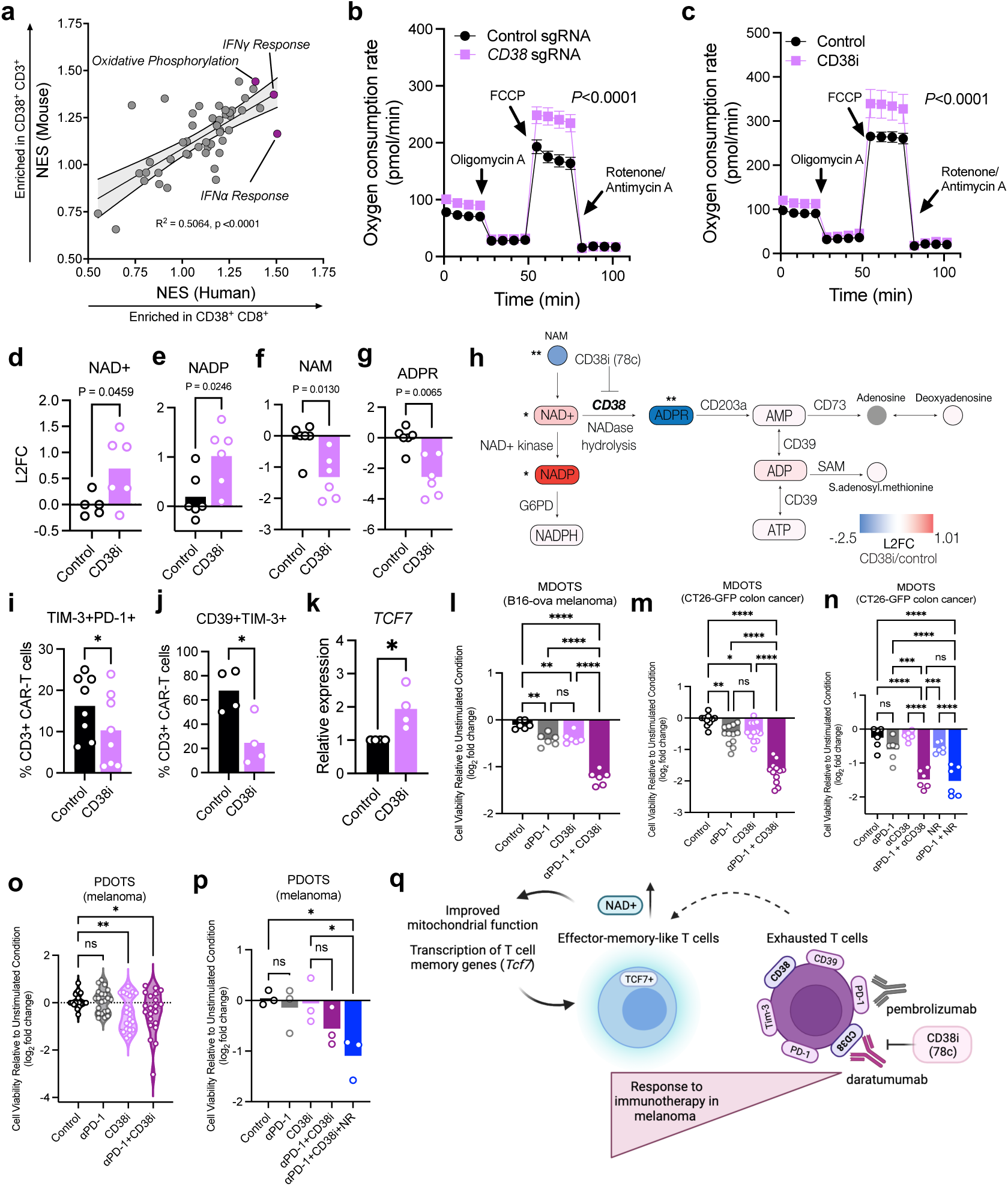
CD38 inhibition restores cellular NAD*^+^* and sensitivity to ICB. **a**, correlation analysis between the GSEA of CD38^+/-^CD8^+^ T cells in human melanoma^10^ and CD38^+/-^CD3^+^ T cells in B16-ova murine melanoma^21^. **b-c,** Oxygen consumption rate (OCR) under basal condition in response to indicated mitochondrial inhibitors of chronically stimulated B7-H3.CAR-T cells form indicated groups. (n=5 biological replicates, 2 independent experiments; 2-way ANOVA with Sidak correction for multiple comparisons). **d-g**, Relative levels of **(d)** NAD^+^ (Nicotinamide adenine dinucleotide), **(e)** NADP^+^ (Nicotinamide adenine dinucleotide phosphate), **(f)** NAM (nicotinamide) and **(g)** ADPR (Adenosine diphosphate ribose) in B7-H3.CAR-T cells in indicated groups (n=6 biological replicates; 2 independent experiments; 2-sided unpaired *t*-test). **h**, scheme of NAD^+^ metabolism and log2 fold change (L2FC) of indicated analytes. **i-j**, surface staining of (**i**) TIM-3^+^PD-1^+^ and (**j**) CD39^+^TIM-3^+^ in B7-H3.CAR-T cells in indicated groups. Means (bars) and individual values (open circles) are shown (n≥4; 2-sided paired *t*-test). **k**, *TCF7* expression in B7-H3.CAR-T cells in indicated groups. Means (bars) and individual values (open circles) are shown (*n*=4; 2-sided unpaired *t*-test). **l-m**, Viability assessment of (**l**) B16-ova MDOTS (n=6 biological replicates, 2 independent experiments), (**m**) CT26-GFP MDOTS (n=12 biological replicates, 4 independent experiments), (**n**) CT26-GFP MDOTS (n=6 biological replicates, 2 independent experiments) with indicated treatments. Means (bars) and individual values (open circles) are shown (one-way ANOVA with Tukey correction for multiple comparisons). **o,** Viability assessment of melanoma PDOTS with indicated treatments. (*n*=21, 7 independent specimens. Mean values (bars) and individual values (open circles) are shown.; mixed effects one-way ANOVA with Geisser-Greenhouse correction and Holm-Sidak correction for multiple comparisons test. **p,** Viability assessment of melanoma PDOTS treated with indicated treatments (*n*=3 biological replicates). Means (bars) and individual values (open circles) are shown; one-way ANOVA with Tukey correction for multiple comparisons. \**P* < 0.05, \*\**P* < 0.01; \*\*\**P* < 0.001, *ns*, not significant. **q,** Scheme demonstrating the effect of targeting CD38 on ICB response by increasing NAD^+^ and *TCF7* expression along with restoring mitochondrial bioenergetics.

As NAD^+^ boosting has been shown to restore mitochondrial fitness and improve T cell function^29^, we next sought to determine if disrupting CD38 could improve mitochondrial metabolic fitness by restoring cellular NAD^+^ levels. Genetic deletion and pharmacologic inhibition of CD38 (78c, CD38i)^30,31^ in T cells increased NAD(H) levels (**Extended Data Fig. 8a-b)**, which was further enhanced with nicotinamide riboside (NR) supplementation (**Extended Data Fig. 8b)**.

## Disrupting CD38 restores NAD^+^ in exhausted T cells

CD38 is a type II membrane glycoprotein whose catalytic domain faces the extracellular space, and despite a well described ‘topological paradox,’ regulates both extra- and intracellular NAD^+^ levels^32^. To define the broader metabolic consequences of targeting CD38 in T cells, we first performed intracellular metabolite profiling of B7-H3.CAR-T cells +/- chronic TCR stimulation using liquid chromatography-mass spectrometry (LC-MS). Chronic TCR stimulation was associated with relative increases in a number of metabolites, including intermediate chain acylcarnitines, glycerol-3-phosphate shuttle metabolic intermediates (glycerol-3-phosphate, dihydroxyacetone phosphate), malate-aspartate shuttle intermediates (malate, aspartate), and NADPH (**Extended Data Fig. 8c-d, Supplementary Table 8**). Of note, NADPH can be generated from NAD^+^ via phosphorylation by NAD^+^ kinase to NADP^+^ and subsequent reduction of NADP^+^ to NADPH by glucose-6-phosphate dehydrogenase (G6PD), an enzyme also enriched in exhausted CD8^+^ T cells in melanoma^33^. Further metabolomic analysis of CD38i-treated, chronically TCR-stimulated B7-H3.CAR-T cells demonstrated increased levels of both NAD^+^ and NADP (**Fig. 4d-e, Supplementary Table 9)**. Thus, high levels of CD38 in exhausted T cells might contribute to reduction in NAD^+^ and NADP levels, the precursors of NADPH, an important metabolite in maintenance of reductive balance and mitochondrial function^34^.

Beyond consuming NAD^+^, CD38 is thought to promote immune suppression via upregulation of adenosine, a potent immune suppressive molecule^35^. Upon CD38 inhibition, we observed decreased levels of the NAD^+^ precursor nicotinamide (NAM) as well as ADP-ribose (ADPR)^36^ (**Fig. 4f-g**). ADPR, the product of NAD^+^ catabolism by CD38, is further catabolized to adenosine by the sequential action of CD203a and CD73^37^ (**Fig. 4h**). While adenosine was not reliably detected in our analysis, related metabolites (e.g, AMP, ATP, ADP, S-adenosyl-L-methionine (SAM) and deoxyadenosine) were detectable and were not significantly altered upon CD38i treatment (**Extended Data Fig. 8e-i**). It is, therefore, possible that other ectoenzymes such as CD73, CD203a, and CD39 expressed on chronically stimulated CAR-T cells (**Extended Data Fig. 8j-k)** compensate for the inhibition of CD38 and contribute to the maintenance of these metabolites^37^. Further, modulation of adenosine levels might depend on T cell extrinsic processes involving multiple cell types in the tumour microenvironment^11,37,38^. LC-MS evaluation of conditioned media of control and CD38i-treated CAR-T cells failed to detect extracellular NAD(P)^+^, and no significant changes in extracellular levels of ADPR, NAM, SAM or deoxyadenosine were observed (**Extended Data Fig. 8l-o, Supplementary Table 10)**, consistent with prior reports that CD38 primarily governs intracellular NAD^+^ levels^39^. These metabolic changes were accompanied by decreased proportions of PD-1^+^TIM-3^+^ and CD39^+^TIM-3^+^ B7-H3.CAR-T cells and increased *TCF7* expression (**Fig. 4i-k**), indicating a less exhausted T cell state, accompanied by elevation of cellular NAD^+^ levels^40^.

## Restoring NAD^+^ to overcome ICB resistance

CD38 blockade with daratumumab partially inhibits CD38 enzymatic activity, but also results in internalisation of the enzyme^41^. To determine if pharmacological inhibition of CD38 and restoration of cellular NAD^+^ pools was sufficient to restore sensitivity to PD-1 blockade, we performed MDOTS profiling of explanted B16-ova and CT26-GFP tumours. CD38i exhibited modest single-agent activity, but significantly increased the response to PD-1 blockade in both B16-ova and CT26-GFP MDOTS (**Fig. 4l-m, Extended Data Fig. 8p**). Consistent with this observation, nicotinamide riboside (NR) supplementation (mimicking the effect of CD38i) enhanced response to PD-1 blockade to a similar extent as disrupting CD38 in CT26-GFP MDOTS (**Fig. 4n**). Further, NAD^+^ depletion by the nicotinamide phosphoribosyltransferase (NAMPT) inhibitor FK866^42^ blunted the effect of dual PD-1/CD38 blockade (**Extended Data Fig. 8q**), confirming a key role for the CD38-NAD^+^ axis in mediating response to ICB. To determine if CD38i would exhibit comparable activity as anti-CD38 (daratumumab) in human tumours, we next evaluated CD38i +/- PD-1 blockade in melanoma PDOTS. CD38i treatment sensitised ICB-resistant melanoma PDOTS to PD-1 blockade (**Fig. 4o, Supplementary Table 11**), an effect that could be enhanced with NR supplementation (**Fig. 4p, Supplementary Table 11**). Taken together, these findings demonstrate the strong therapeutic potential of targeting CD38 to overcome ICB resistance, and underscore the importance of NAD^+^ as a vital metabolite for maintaining and restoring T cell function (**Fig. 4q**).

## DISCUSSION

Here we show that CD38^+^CD8^+^ T cells are associated with ICB resistance and exhibit features of T cell exhaustion, including reduced proliferative capacity, decreased effector function, upregulation of co-inhibitory receptors, and altered mitochondrial bioenergetics. Genetic deletion or pharmacologic inhibition of CD38 restored T cell proliferation and cytotoxic function in association with increased NAD^+^ levels and increased *TCF7* expression, and disrupting CD38 restored sensitivity to ICB in murine- and patient-derived *ex vivo* tumour models. Via its capacity to regulate intracellular NAD^+^ levels, CD38 influences several cellular processes including mitochondrial function^14,29^, metabolism, and gene transcription^36^. Indeed, CD38 inhibition restored NAD(P)^+^ pools alongside improved mitochondrial function and increased expression of *TCF7*, resulting in improved T cell effector functions and improved response to ICB in a NAD^+^ dependent manner. While SIRT1, a member of the NAD^+^-dependent sirtuin family of deacetylases, has been shown to regulate T cell memory gene expression in murine CD4^+^ T cells ^17^, the precise molecular mechanism(s) whereby modulation of NAD^+^ metabolism regulates TCF7 expression and T cell effector-memory state in tumour-infiltrating CD8^+^ T cells has yet to be defined.

The mechanism whereby CD38 is transcriptionally regulated during chronic antigen exposure and subsequent T cell exhaustion in cancer also remains incompletely understood. Similar to *PDCD1* (encoding PD-1) and *ENTPD1* (encoding CD39), the *CD38* locus is open in exhausted T cells in cancer and chronic viral infection^13,43^. Interestingly, studies in patients with hepatitis C virus (HCV) confirmed high expression of *CD38* in the setting of chronic viral infection, and demonstrated normalisation of *CD38* expression following effective antiviral therapy^44,45^, in contrast to *ENTPD1* and *PDCD1* whose elevated expression is maintained even after successful viral clearance, suggesting permanent epigenetic scarring^40^. Thus, despite the strong co-expression of CD38 with *PDCD1* (PD-1), *ENTPD1* (CD39), and other putative exhaustion markers, the factors governing up- and -downregulation of CD38 and extent of reversibility will require further investigation.

Resistance to ICB cancer immunotherapy in melanoma and other cancers remains a significant clinical challenge with available second line combination therapies benefiting a minority of patients^6^. The observed efficacy of dual PD-1/CD38 blockade, particularly in patient-derived *ex vivo* tumour models from patients with clinical ICB resistance, supports further preclinical and clinical development of CD38-directed therapeutic strategies to overcome ICB resistance. As CD38 monoclonal antibodies (e.g., daratumumab) are already FDA-approved for other indications (e.g., multiple myeloma), there is a clear path to rapid translation of these findings to clinical trials to combat ICB resistance.

## Data availability

The datasets generated and analysed in this study are available upon request. Metabolomics raw data will be available on Metabolomics Workbench (a metabolomics data depository).

## Code availability

The codes used to figure generation and data analysis in this study are available upon request.

## METHODS

### Patient samples

Tumour samples were collected and analysed according to Dana-Farber/Harvard Cancer Center IRB-approved protocols (IRB protocols: 11-181, 02-240). A cohort of patients (Supplementary Table 5 and 11) treated at Massachusetts General Hospital was processed for PDOTS profiling and patient derived cell lines and TILs were isolated and expanded (see relevant sections). These studies were conducted according to the Declaration of Helsinki and approved by the DF/HCC IRB Response to treatment was determined radiographically, as previously described^21^.

### Analysis of human scRNAseq data

*<u>Single-cell RNA-seq datasets and pre-processing:</u>* we used two scRNA-seq datasets ^10,18^ containing tumour biopsies of cancer patients treated with checkpoint immunotherapy. These datasets were pre processed and analysed using Scanpy ^46^ version 1.9.4. Expression levels for the single smart-seq2 dataset that we have previously published ^10^ were quantified as transcripts per million (TPM) and were log transformed as follows: *log*_2_ (*TPM* + 1). The NSCLC dataset was droplet-based and contained read counts. In this dataset, we filtered out cells having more than 10% of mitochondrial genes count and removed cells expressing less than 200 genes. Genes that were expressed in less than 3 cells were omitted. We then applied Scrublet ^47^ and removed cells having “DoubletScore” higher than 0.3. Expression levels were then normalised to a standard target sum of 10,000 reads per cell, and log transformed as follows: *log*_2_ (*normalised counts* + 1). For the smart-seq2 dataset, annotations for cellular clusters, including t-distributed stochastic neighbour embedding (tSNE) ^48^ for dimensionality reduction, as well as genes passing QC, were used according to the published data provided by the authors: (https://singlecell.broadinstitute.org/single_cell/study/SCP398/defining-t-cell-states-associated-with-response-to-checkpoint-immunotherapy-in-melanoma) ^10^.

*<u>Identification of CD8^+^ T cells:</u>* in the human melanoma dataset, CD8^+^ classification was performed as was previously described ^10^. In NSCLC dataset^18^, we used a stringent criteria such that a CD8^+^ T cell must express *PTPRC* (*CD45*), *CD3E* and *CD8A* or *CD8B*, and cannot express either *NCR1*, *NCAM1* or *FOXP3* ^10^. A gene was considered as “expressed” if *log*_2_ (*normalised counts* + 1) > 1, and “not expressed” otherwise. CD8^+^ T cells were extracted out of all of the sequenced cells passing our quality control, as they went through CD3^+^ positive cell filtration by flow cytometry.

*<u>Assessing the predictive value of CD38 for cancer patient response:</u>* in each dataset, we focused only on CD8^+^ T cells as previously described (identification of CD8+ T cells section above) and divided them into two groups according to their expression of *CD38*. For the melanoma dataset, *CD38* considered to be expressed when the *log*_2_(*TPM* + 1) was larger than 2.5. For the NSCLC dataset, *CD38* considered to be expressed when the *log*_2_(*normalised counts* + 1) was larger than 1. We computed a score per sample by taking the ratio between the number of cells classified as CD38^+^ and those that were CD38^-^. We calculated the Receiver Operating Curve (ROC) and the Area Under the Curve (AUC) score and applied a two-sided unpaired *t*-test to calculate the P-value for the comparison of the ratios between responders and non-responders.

*<u>Differential expression analysis:</u>* in order to identify differentially expressed genes between CD38^+^ and CD38^-^ CD8^+^ T cells, we first considered only genes that are expressed in more than 10% of the cells, in at least one of the groups. Following, for each gene we count the number of cells that express it with an expression level (*log*_2_(*TPM* + 1) > 2. 5) or (*log*_2_(*TPM* + 1) ≤ 2. 5). We then applied Fisher’s Exact test for the corresponding contingency table. Significant genes were those with an adjusted P-value < 0.05 after correcting for multiple hypotheses using the Benjamini-Hochberg false discovery rate^49^ and *log*_2_(*FoldChange*) > 0. 5. The set of differentially expressed genes in each group for the melanoma dataset ^10^ (CD38^+^/CD38^-^) is listed in Supplementary Table 3.

*<u>Gene set enrichment analysis (GSEA):</u>* we conducted GSEA for the melanoma dataset^10^ using the “GSEApy” library version 1.0.4 in python^50^. First, the dataset was subset to only include protein coding genes passing QC according to Sade-Feldman et al.^10^. We then filtered out genes having mean expression of zero in at least one of the two groups (CD38^+^/CD38^-^). Differentially expressed genes between groups (CD38^+^/CD38^-^) (Supplementary Table 6) were ranked as follows: − *log*_10_(*pvalue*) ∗ *log*_2_(*FoldChange*). Ranked lists of differential genes were created using signed p-values and passed to *GSEA Preranked* to search for enriched gene sets^51^. The Hallmark TNFα Signalling via NFκB had a negative running enrichment score and was excluded from Extended Data Fig. 7a and Fig. 4a for visual clarity when compared to the B16-ova dataset (see below).

*<u>Analysis of CD38^+^CD8^+^ T cells in patients treated with single PD-1 blockade or dual PD-1 and CTLA-4</u> <u>blockade:</u>* we subset the data to include only CD8+ T cells. Quality-control, filtering, and normalisation were performed as previously described^10^. Additional analysis and plotting were performed using Scanpy (v.1.9.3) and Scipy (v.1.10.1). Two violin plots showing CD38 expression were generated with separation by immune checkpoint blockade target, patient response status, and time at which the sample was taken (pre- and post-therapy). The expression values of *CD38* are after each cell transcriptome was scaled to sum to 100,000, and expression values were further normalised with log1p, finally obtaining log[TP100K + 1] values for each gene. We used the nonparametric 2 sample Kolmogorov-Smirnov test to evaluate whether the distribution of CD38 was the same comparing non-responder patients to responders. We conducted this test four times to compare within ICB treatment and time the sample was taken. Significant differences were those with a p-value < 0.005.

### Analysis of B16 scRNAseq

(Gene Expression Omnibus (GEO): GSE225713). Processed scRNA-seq data was obtained from a previously published dataset ^20^. We subset to only include CD8+ T cells from tumours, draining and non-draining lymph nodes (dLN and ndLN respectively) at day 7, day 10, and day 16 after B16F10 melanoma inoculation. Quality-control, filtering, and normalisation were performed as previously described ^20^. Additional analysis and plotting were performed using Scanpy (v. 1.9.3). Principal component analysis (PCA) and nearest neighbour graphs were calculated to visualise on a UMAP plot. Dot plots depicting mean expression and fraction of cells in groups expressing the gene are shown. The first plot of *Cd38*, normalised to values between 0 and 1, with grouping by tissue of origin. We then plotted the expression of *Cd38* and *Tcf7*, normalised in the same manner as before, with grouping by the tissue of origin and the time point of collection. Using the original UMAP representation generated by Bod et al ^20^, we plotted cell type, tissue of origin, and *Cd38* expression. The expression values of *Cd38* are after each cell transcriptome was scaled to sum to 10,000, and expression values were further normalised with log1p, finally obtaining log[TP10K + 1] values for each gene.

### Analysis of B16-ova scRNAseq

scRNA-seq data for CD45+ tumour-infiltrating immune cells isolated from C57BL/6J mice inoculated with subcutaneous B16-ova tumours treated with ICB were previously described ^21^ (Gene Expression Omnibus (GEO): GSE217160). In this analysis, data were subset to only include cells isolated from control (vehicle/IgG-treated) and anti-PD-1 (vehicle/anti-PD-1-treated) groups. Quality-control, filtering, and normalisation were performed as previously described ^21^. Additional analysis and plotting were performed using Scanpy (v.1.9.3). Uniform Manifold Approximation and Projection (UMAP) plots were generated using the built-in Scanpy Principal Component Analysis (PCA) and nearest neighbour graph functions. PCA embeddings were corrected to minimise technical batch effects between experiments using Harmony ^52^. Cells (n=19,214) were then grouped into 26 distinct clusters using the Leiden algorithm. Galaxy plots depicting cluster density and tracksplots are downsampled to match the treatment group with the smallest cell count (control, n=6,638). To gain more granularity between the T and NK cell subtypes, sub-clustering was performed on cells in specific clusters expressing *Cd8a, Cd4* and *Ncr1* transcripts based on the built-in Scanpy function one-versus-rest differential expression. After reclustering, new PCA and UMAP embeddings, nearest neighbourhood graphs, and Harmony batch corrections were calculated on a newly calculated set of 10,000 highly variable genes. T and NK cells (n=5,464) were then grouped into 8 distinct clusters using the Leiden algorithm. Galaxy plots depicting cluster density and tracksplots are downsampled to match the treatment group with the smallest cell count (control, n=1,271).

<u>*Differential gene expression:*</u> to assess differential gene expression between T cells clusters based on CD38 expression, cells in clusters expressing Cd8a, Cd4, and Ncr1 transcripts (as described above) were further categorised based on the built-in Scanpy function Score Genes using Cd38 transcript expression to calculate a CD38 expression score for each cell. Cells were then assigned to one of two groups (CD38^+^, score > 0 or CD38^-^, score <= 0). Differentially expressed genes between CD38^+^ and CD38^-^ T cells (Early Effector CD8+, Progenitor CD8+, Terminal Effector CD8+, *Mki67*+, and CD4+) were calculated using a logistic regression model ^53^.

*<u>Gene set Enrichment analysis:</u>* to perform GSEA analysis, ranked lists of differential genes were created using signed p-values calculated by the logistic regression model and passed to *GSEA Preranked* using “GSEApy” library version 1.0.4 in python ^50^ to search for enriched gene sets ^51^. (Supplementary Table 7).

### scRNA seq analysis of human melanoma validation cohort

scRNA-seq data for CD8^+^ tumour-infiltrating immune cells isolated from fresh metastatic melanoma samples (n = 39, Supplementary Table 1) were collected from Massachusetts General Hospital in collaboration with Dr. Genevieve Boland and sequenced via 10x Genomics chromium controller and 3’ transcriptomics reagent kits to form our scRNA-seq atlas. Processing and quality control steps were performed, as described here (https://hdl.handle.net/1721.1/144938). Libraries were merged together and principal component analysis (PCA) was performed, using the genes detected in all samples. After obtaining the low-dimensional embeddings, we performed Uniform Manifold Approximation and Projection (UMAP) on the top 40 principal components to visualise the dataset. We identified CD8^+^ T lymphocytes (*CD3E*, *CD8A*, *GZMA*) from the cohort using known cell-type specific markers. To identify the proliferating cells, we used the CellCycleScoring function from Seurat to calculate the S phase score and G2M phase score for each cell ^54^. Proliferating cells were not used in the CD8^+^ T-cell specific analyses. In the CD8^+^ T compartment, we used Harmony to integrate cells from 35 samples that contained CD8 T cells and identified clusters through the Louvain clustering algorithm. After checking the quality of the cells, we found a cluster with high mitochondrial percentage and number of gene features, which are likely doublet cells. With the doublet cells removed, 22,008 cells in 10 clusters remained in the study, indicated in (Extended Data Fig. 1l). Percentage of CD8^+^ T-cells expressing CD38 was analysed, and the proportion in ICB responsive vs. non-responsive was determined. For the assessment of the proportion of CD38^+^ CD8 T cells ROUT method for removing outliers was applied. UMAPs visualising *CD38* and *TCF7* expression across CD8^+^ T-cell subtypes were created by normalising cell counts to (to sum to 1,000) and logarithmized. Max cutoff for UMAP normalised gene expression was set at 3.6. For co-expression deviation proportion (CDP), we performed pairwise comparisons between pairs of naïve and exhaustion markers to detect their co-expression patterns in CD8^+^ T-cells. For each gene pair, we found the proportion *a* of CD8^+^ T cells expressing the first gene and the proportion *b* of CD8^+^ T cells expressing the second gene. Assuming random distributions of each gene across the population, we expect some proportion *e = ab* to express both genes of the gene pair. Then, we found the actual proportion of gene expression in the gene pair. We calculated what we termed the co-expression deviation proportion (CDP) as follows: *(f - e) / e*. In the case that *e = 0*, if also *f = 0*, then we set CDP for that gene pair to be *0*; else the CDP for that gene pair is *1*.

### CyTOF immune profiling

*<u>Mass cytometry panel design and heavy-metal conjugation of antibodies</u>*. Some antibodies were obtained pre-conjugated to heavy-metal isotopes from Standard BioTools (formerly Fluidigm). In-house conjugations were performed as needed using the MaxPar X8 or MaxPar MCP9 antibody-labelling kits (Standard BioTools) following an optimised and updated protocol^55^ and according to manufacturer instructions. The allocation of targets to appropriate heavy-metal isotopes was based on the sensitivity of the mass cytometer (e.g., lower abundance targets were placed on higher sensitivity channels) and to avoid potential signal spillover as previously described^56^. *<u>Sample</u> <u>preparation</u>*. Cryopreserved PBMC samples (Supplementary Table 2) were thawed, washed, and resuspended in PBS + 5mM EDTA before cisplatin staining (Sigma) for 1 min at RT. Cells were washed and fixed with 1.6% PFA in PEB (PBS + 5mM EDTA + 0.5% BSA) for 10 min at RT. Cells were washed twice with PEB and stored at -80°C until staining. *<u>Sample barcoding</u>*. To minimise batch-to-batch variation and improve data quality, samples were barcoded prior to antibody staining using a 20-Plex Pd Barcoding Kit (Standard Biotools) following manufacturer instructions. Briefly, after counting, 1 million cells were washed with MaxPar Barcode Perm Buffer (Standard Biotools) twice and barcoded with unique combinations of Pd isotopes for 30 min at room temperature on a shaker in Maxpar Barcode Perm Buffer. Cells were washed twice with eBioscience perm buffer and once with Maxpar Cell Staining Buffer (CSB, Standard BioTools) and pooled into a single tube. *<u>Antibody staining</u>*. Antibody cocktail was prepared prior to staining according to previously determined antibody titers at 25 μL of staining volume per 1 million cells. The antibody cocktail was aliquoted and stored at -80°C until staining. The pooled barcoded sample was stained for 10 minutes at room temperature with Fc receptor blocking solution (BioLegend) followed by 1h incubation with the antibody staining cocktail at room temprature. PBMCs were stained using a panel of 43 antibodies (Supplementary Table 12). Cells were washed twice with CSB and resuspended in intercalation solution (4% PFA in PBS and 0.5 mM iridium intercalator (Standard BioTools) for 20 min at RT. Stained samples were pelleted and frozen at -80°C until acquisition. *<u>Data acquisition</u>*. Before acquisition, samples were washed once in CSB and twice in CAS (Standard BioTools) and filtered through a cell strainer (Falcon). Cells were then resuspended at 0.7 million cells/mL in CAS supplemented with EQ4 element calibration beads (Standard Biotools) and acquired on a Helios mass cytometer (Standard BioTools). All samples met quality control standards. Manual gating of FCS files was performed using CellEngine (CellCarta, Montreal, Canada). *<u>Data analysis</u>*. PBMCs were separated into different cell populations as shown in Supplementary Data Fig. 3. Abundances of immune cell subsets between patients with clinical response (CR, PR, SD for more than 6 months) and patients without clinical response (PD, SD for less than 6 months) were compared.

### Plasma collection and OLINK plasma proteomics analysis

Olink profiling of blood samples (collected using DF/HCC IRB approved Protocol 11-181) from patients with metastatic melanoma at MGH was performed as previously described^21^. Olink Proximity Extension Assay (PEA) for high-multiplex analysis of proteins was performed as previously described^57^. The full library (Olink ® Explore 1536) consists of 1472 proteins and 48 control assays, divided into four 384-plex panels. Four overlapping assays of IL-6, IL-8 (CXCL8), and TNF are included for quality control (QC) purposes. In the immune reaction, 2.8 mL of sample is mixed with PEA probes and incubated overnight at 4°C. Olink’s relative protein quantification unit on a log2 scale and values are calculated from the number of matched counts on the NovaSeq run. Data generation of normalised protein expression (NPX) consists of normalisation to the extension control (known standard), log2-transformation, and level adjustment using the plate control (plasma sample).

### CAR-T cell generation

B7-H3.CAR-T were generated as described previously^23^. Briefly, the B7-H3 376.96 mAb scFv was included into a CAR construct containing a human CD8α hinge and transmembrane domain, CD28 intracellular costimulatory domain and CD3ζ intracellular signalling domain. The B7-H3.CAR cassette was cloned into the retroviral vector SFG and provided to us by Dr. Gianpiettro Dotti (University of North Carolina at Chapel Hill, Chapel Hill, NC). Retroviral supernatant was generated by transfecting 293T cells with a plasmid mixture (the B7-H3.CAR retroviral vector, the Peg-Pam-e plasmid encoding MoMLV gag-pol, and the RDF plasmid encoding the RD114 envelope) using the GeneJuice transfection reagent (Merck Millipore). Viral supernatant was collected 48 and 72 hours after transfection. Peripheral blood mononuclear cells (PBMCs) isolated from whole blood were plated with CD3 (Miltenyi, 130-093-377) and CD28 (BD Bioscience, 556620) mAbs. Activated T cells were transduced with the retroviral supernatant using retronectin-coated plates (Takara Bio Inc, T100B). T cells were harvested after three days and expanded in complete medium (45% RPMI-1640 (Corning, 10-040-CM) 45% Click’s medium (Irvine Scientific, 9195-500ML), 10% FBS (Sigma-Aldrich, F2442), 100 unit/mL of Penicillin and 100 mg/mL of streptomycin (Sigma-Aldrich, P4458) with IL-7 (10 ng/mL; PeproTech, 200-07) and IL-15 (5 ng/mL; PeproTech, 200-15). CAR transduction efficiency was assessed by flow cytometry using the FITC-labelled human B7-H3 (4Ig) protein (Acro Biosystems, B7B-HF2E7). B7-H3.CAR-T cell products were derived from CD3^+^ PBMCs from different human normal donors^23^, and contained a mixture of CD8^+^, CD4^+^ T cells, and a small portion of other CD3^+^ cells (Supplementary Fig. 4a).

### Cell culture

CAR-T cells were cultured in (45% RPMI-1640 (Corning) 45% Click’s medium (Irvine Scientific), 10% FBS (Sigma-Aldrich), 100 unit/mL of Penicillin and 100 mg/mL of streptomycin (Thermo Scientific, 15140122) with IL-7 (10 ng/mL; PeproTech) and IL-15 (5 ng/mL; PeproTech), supplemented by 10% BenchMark™ Fetal Bovine Serum (GeminiBio, 100-106), or ImmunoCult (STEMCELL, 10981) + 50 ng/mL IL-2 (PeproTech, 200-02). When indicated CAR-T were treated with CD38 inhibitor 1uM 78c (6391, Toris). CD8^+^ TILs were cultured in ImmunoCult (STEMCELL, 10981) + 50 ng/mL IL-2 (PeproTech, 200-02). Jurkat and patient derived tumour cell lines 10101 and 10164 were cultured in RPMI (Corning, 10-040-CV) supplemented with 10% BenchMark™ Fetal Bovine Serum (GeminiBio, 100-106) and 1% penicillin-streptomycin (Thermo Scientific, 15140122). Patient derived cell line 10170 were cultured in SMGM (Lonza, CC-3182). All cells were cultured at 37 °C in 5% CO2 humidity and were tested routinely for mycoplasma using MycoAlert mycoplasma detection kit (Lonza, LT07-318) and appropriate positive control (Lonza, LT-07-518).

### *In vitro* T cell chronic stimulation

For chronic stimulation assays, we modified previously developed protocols^29,58^. B7-H3.CAR-T cells or patient derived CD8^+^ expanded TILs were activated with washed aCD3/aCD28 dynabeads (ThermoFisher Scientific, 11161D), according to the manufacturer protocols, in a 1:1 ratio of beads to cells, 20 ng/mL of IL-2 (PeproTech, 200-02), 0.6ug/ml aFAS neutralising antibody (EMD Millipore, 05-338) and RPMI (Corning, 10-040-CV), supplemented with 10% FBS and 1% penicillin/streptomycin. After 3 days, the beads were removed using a magnet and the cells were divided into acute and chronic conditions. Acute and chronic were cultured in the media above, and only the chronic got restimulated with 1:1 aCD3/aCD28 dynabeads to T cells ratio. Media and beads were replaced every 2-3 days for a total of 10-14 days.

### Generation of CRISPR-edited T cells

For *CD38* deletion in B7-H3.CAR-T cells, 1×10^6^ T cells were transfected with Cas9/RNPs complexes of CD38 targeting guide RNA 5’-AGUGUAUGGGAUGCUUUCAA -3’ (Synthego) using Lonza 4D-Nucleofector™ X Unit (Lonza, AAF-1003X). Transfection was carried out in P3 Primary Cell 4D-Nucleofector™ Solution (Lonza, V4XP-3032) and program E0-115 was used. Immediately after transfection, pre-warmed, incubator equilibrated ImmunoCult media + IL-2 (50 ng/mL) was added to the T cells, followed by culturing for 3 days. CD38 Deletion efficiency was tested 3-6 days post transfection using flow cytometry. Other guides were tested that yielded lower editing efficiency (Supplementary Data Fig. 4b, Supplementary Table 13.

### Flow cytometry and cell sorting

For both flow cytometry and cell sorting, Human TrueStain FcX (Biolegend, 422302) or mouse FcR blocker (Miltenyi 130-092-575) were used for blocking Fc receptors before labelling cells. To discriminate between live and dead cells, we used Zombie Violet Dye (Biolegend, 423114) for 15 min at 4°C, followed by surface labelling of cells for 30 min at 4°C, using standard protocols^10^. The antibodies used for cell surface labelling can be found in Supplementary Table 14. Intracellular staining for human TCF7 was performed following surface staining, fixation and permeabilization using eBioscience™ Intracellular Fixation & Permeabilization Buffer Set (Thermo Fisher, 88-8824-00), according to the manufacturer’s instructions. Flow data was acquired with the multi-spectral Cytek Northern Lights instrument and MA900 Sony Cell sorter and when cell sorting was performed with MA900 Sony Cell sorter. Analysis was done using FlowJo software version 10.8.1. Sorting gating strategy (Supplementary Data Fig. 4c-d). Following sorting, post sorting validation flow was performed. Flow cytometry gating strategy (Supplementary Data Fig. 4-8). List of antibodies that were used can be found in Supplementary Table 14.

### RNA Isolation and RT-qPCR

Total RNA was extracted from B7.H3 CAR-T cells using the RNeasy Kit (Qiagen, 74104) according to the manufacturer’s protocol. RNA levels were quantified using the Blaze Taq one-step SYBR Green RT-qPCR kit (GeneCopoeia, QP070) according to the manufacturer’s protocol and acquired on the ROCHE Lightcycler-96 system. Primer sequences can be found in Supplementary Table 15. RNA samples were normalised to *GAPDH* or *TUBB* and the 2^-ΔΔCt^ was calculated to obtain the relative gene expression.

### Organotypic tumor spheroid preparation and microfluidic device culture

PDOTS and MDOTS were generated as previously described ^24^. In brief, fresh tumour specimens from mice or humans were received in full DMEM media (Corning, 10-013-CV), or RPMI (Corning, 11875-093) for CT26 tumours supplemented with 10% BenchMark™ Fetal Bovine Serum (GeminiBio, 100-106) and 1% penicillin-streptomycin (Thermo Scientific, 15140122) on ice and minced in a 10 cm plate using sterile forceps and scalpels, if samples arrived late, they were stored in tissue storage media (Miltenyi, 130-100-008) and processed the day after. Minced tumours were resuspended in full DMEM (RPMI for CT26 tumours) and were passed over 100-μm and 40-μm filters sequentially to generate the S1 (>100 μm), S2 (40-100 μm), and S3 (<40 μm) fractions. S2 and S3 were washed off the filter with fresh full media and were rested in ultra low attachment plates in (Corning, 3471) in the incubator until loading into the device. S1 fraction was washed from the filter using fresh full media supplemented with 100 U/mL collagenase type IV (Life Technologies, 17104019), and 15 mM HEPES (Gibco, 1560-080). S1 with collagenase was incubated for 15-30 minutes at 37°C followed by the addition of an equal volume of media and subsequent filtering. The S2 fraction was pelleted and resuspended in type 1 rat tail collagen (Corning, 354236) at a concentration of 2.5 mg/mL with 10x PBS + phenol red (Sigma-Aldrich, 114537-5g). A pH of 7.0-7.5 was confirmed with PANPEHA Whatman paper (Sigma-Aldrich, 2629990) after titrating the solution with NaOH (Sigma-Aldrich, 1.09138.1000). The spheroid-collagen mixture (10 μL) was added into the centre channel of the AIM 3D microfluidic device (DAX-01, AIM Biotech). Collagen hydrogels with PDOTS/MDOTS were incubated for 20 minutes at 37°C in sterile humidity chambers before being hydrated with media in the side channels with or without treatments. PDOTS were treated with anti-PD-1 (250 μg/mL, pembrolizumab), anti-CD38 (200 μg/mL, daratumumab), anti-PD-1+anti-CD38 combination, 1μM 78c (6391, Toris), combined anti-PD-1 + 78c, combined anti-PD-1 + 78c+ NR (Sigma-Aldrich, SMB00907-50MG) or left untreated. MDOTS were treated with isotype control IgG2a (Bio X Cell, 2A3, 10 μg/mL), anti-PD-1 (Bio X Cell, RMP1-14, BE0146, 10 μg/mL or Biolegend, 114102, 10 μg/mL), anti-CD38 (Novus, NIMR-5, 0.5, 1.0, 5.0 μg/mL), 78c (0.5, 1.0, 5.0 μM), 100 μM of nicotinamide riboside chloride (NR, Sigma-Aldrich, SMB00907-50MG), anti-CD8β (Bio X Cell, BE0223) and 1 nM of FK866 NAMPT inhibitor (Tocris, 4808). S1 and S3 were banked using BAMBANKER freezing media (Bulldog Bio, BB02) and kept for downstream analysis by flow cytometry (see section) or patient derived tumour cell line with matched TILs formation (see relevant section below).

### Generation of murine tumours for MDOTS and TILs

The designs of animal studies and procedures were approved by the Charles River Laboratories IACUC committees. Ethical compliance with IACUC protocols and institute standards was maintained. Murine pathogen testing and mycoplasma testing was performed prior to tumour inoculations. B16-ova MDOTS were prepared from tumours using wild-type female C57BL/6J mice (7-8 weeks old, Charles River Laboratories), 1.0 × 10^6^ B16-ova cells (kindly provided by Dr. Debattama Sen, MGH) were resuspended in sterile Ca- and Mg-free PBS (Gibco) and subcutaneously injected into the flank on day 0. CT26-GFP MDOTS were prepared using wild-type female BALB/c mice (7-8 weeks old, Charles River Laboratories). 2.0 × 10^6^ CT26-GFP cells (Kindly provided Dr. Moshe Sade-Feldman, MGH) were resuspended in sterile Ca- and Mg-free PBS (Gibco) and subcutaneously injected into the flank on day 0. Pre-specified endpoints for tumour size were adhered to as defined by IACUC protocols, including 2.0 cm in maximum dimension for moribund end point. CO_2_ asphyxiation was used to euthanize mice. At least five tumours were collected for each MDOTS experiment. No *in vivo* drug challenges were performed so randomisation and blinding were not performed. Beginning on day 6 after challenge, tumour volumes (TV) were estimated using longest dimension (length) and the longest perpendicular dimension (width), using the formula (*L* × *W*2)/2. Tumour volumes were assessed every 3–4 days. Mice were euthanized 10-14 days after inoculation and tumours were harvested and shipped in DMEM or issue storage media (Miltenyi, 130-100-008) on ice. For CT26 tumour-derived CD8^+^ TILs, frozen single cell suspensions from MDOTS processing (see above) were thawed and CD8^+^ TILs were isolated using Cd8a (LY-2) Microbeads (Miltenyi Biotec, 130-117-044). T cells were cultured in RPMI (Corning, 10-040-CV) supplemented with 10% BenchMark™ Fetal Bovine Serum (GeminiBio, 100-106) and 1% penicillin-streptomycin (Thermo Scientific, 15140122) with 20 ng/mL murine IL-2 (Peprotech, 212-12). Cells were treated with anti-CD38 (Novus, NIMR-5) 1 μg/mL or isotype control IgG2a (Bio X Cell, 2A3, 10 μg/mL) for 3 days and were analysed using flow cytometry.

### PDOTS and MDOTS viability assessment

PDOTS and MDOTS staining and viability analysis was done as previously described ^24^. In brief, staining was done in microfluidic devices by adding acridine orange/ propidium iodide (AO/PI) solution (Nexcelom, CS2-0106), diluted 1:1 in full media and 12ug/ml Hoechst (Invitrogen, H3570) or 12 μg/mL Hoechst Hoechst and 0.8 μg/mL PI alone (Thermo Scientific, P3566). After incubation with fluorescent dyes (30 minutes, 37°C), images were acquired using a Nikon Eclipse NiE fluorescence microscope, using X4 lens in 2 or 3 colours. Images were analysed using the NIS-Elements AR software package and live/dead cell quantification was obtained by measuring the total cell area of acridine orange for live cells, PI for dead cells and Hoechst for total cells. Percent change and log2FC (L2FC) data were generated using raw fluorescence data (live) for given treatments relative to control conditions.

### PDOTS immunofluorescence

For live fluorescence staining, PDOTS side channels culture media was replaced with a blocking solution of 2% BSA (Sigma, A7906) in PBS for 30 min in room temp, followed by staining with fluorescently conjugated antibodies against CD45, CD38, PD-1, CD39, CD8 (See Supplementary Table 14) and 12 μg/mL Hoechst (Invitrogen, H3570) in blocking solution for 45 min at RT in the dark. Staining solution in the side channels was washed twice with PBS + 0.1% Tween 20 (Fisher Scientific, BP337) and replaced with PBS for imaging. Images were acquired using Leica Thunder system in 20X phase lens and 40X water lens and images were digitally cleared using Leica LAS-X THUNDER module.

### Generation of patient derived tumour cell lines and tumour infiltrating lymphocyte cultures

Fresh single cell suspension remaining from PDOTS processing (S3) was grown in SmGM-2 Smooth Muscle Cell Growth Medium (Lonza, CC-3181) supplemented with SmGM-2 SingleQuots Supplement Pack (Lonza, CC4149). After 24 hrs, the supernatant from the culture was removed and set aside to enrich for CD8^+^ TILs. Adherent cells remaining in the culture were expanded until a stable tumour cell line was established. Following expansion and mycoplasma testing (Lonza, LT07-318), tumour lines underwent SNaPshot sequencing. Some of the fast growing tumour cell lines were further grown in RPMI (Corning, 10-040-CV), supplemented with 10% FBS 1% penicillin-streptomycin. T cells were isolated from the supernatant using CD8 MicroBeads (Miltenyi, 130-045-201) according to the manufacturer’s instructions. T cells were activated for 3 days using 3 μg anti-human CD3 and CD28 antibodies (Biolegend, 300438, 377604) and 50 ng/mL of IL-2 (Peprotech, 200-02) in ImmunoCult-XF media (Stemcell Technologies, 10981), and expanded for 2-3 weeks until reaching the appropriate amount of T cells (usually >10 x 10^6^ cells). Media+ IL-2 was refreshed every 2-3 days.

### Multivariate linear regressions analysis

Multivariate linear regressions were performed with R Statistical Software (v4.3.2). Features of interest were stratified into the following groups: flow cytometry panels with CD4^+^ proportions, flow cytometry panels with CD8^+^ proportions, and clinical features of interest. The flow cytometry panel measuring CD45^-^CD38^+^ proportion was included in models for both CD4^+^ panels and CD8^+^ panels. For each set of features, a multiple linear regression was fitted to predict PDOTS response (% change from control) following anti-PD-1 plus anti-CD38 treatment. Visualisation of the slope estimates and p-values for each model was performed with Python programming language (v3.8.16), matplotlib (v3.7.1) and seaborn (v0.12.2). 0.05 was the p-value threshold used to determine significance.

### T cell proliferation assay

For acute versus chronic proliferation measurements total cell number for acute and chronic was counted at day 0 and every 2-3 days until day 10 using Countess automated cell counter (Thermo Fisher). For Incucyte® proliferation assay plastic flat bottom 96 well plates were coated with 5 μg/mL human fibronectin in PBS for 1h in 37°C. Plates were washed 3 times in PBS and dried under sterile conditions. T cells were cultured in triplicates (30,000-50,000 per well) and allowed to adhere for 15 min before imaging. Cells were imaged every 2-4 hours using the Incucyte® Live-Cell Analysis System phase contrast 10X lens for 3 days. Analysis was performed using Incucyte® non adherent Cell-by-Cell Analysis Software Module and the T cells Object Count Per Well, normalised to 0d0h0m was used. Graphs show a summary of 3 biological repeats, with three technical replicates for each sample.

### T cell-tumour cytotoxicity assay

For Incucyte® tumour cytotoxicity assay, tumour cell lines were cultured in their growth media (RPMI or SmGM-2), 1 day before co-culture with T cells to allow cell spreading. On the day of the assay tumour media was replaced with fresh media with Incucyte® Cytotox Green Dye (Sartorius, 4633), according to the manufacturer instructions and incubated for 10 min prior T cells addition. T cells (CAR-T cell or matched CD8^+^ TILs) were added in different E:T ratios, ranging 1:1, 1:2, 1:5 for CAR-T cells and 1:1, 2:1, 5:1 for CD8^+^ TILs. Co-culture was imaged every 2-4 hours using Incucyte® Live-Cell Analysis System phase contrast 10X lens and green fluorescence for 3 days. Analysis was performed using Incucyte® standard Analysis Software Module and the graphs show the GCU metric that represents integrated intensity of the CytoTox dye-labelling dead cells. Graphs show a summary of 3 biological repeats, with three technical replicates for each sample, each normalised to 0d0h0m GUC values.

### Analysis of human tumours and human T cells by flow cytometry

Single cell suspension from PDOTS processing (S3; see PDOTS generation section) was processed for flow cytometry. If frozen, single cell suspension was thawed into ImmunoCult-XF media (Stemcell Technologies) and incubated at 37°C for 1 hour, to allow cell recovery. Tumours were stained with conjugated fluorescent anti-human monoclonal antibodies against CD3, CD4, CD279, CD366(Tim-3), CD38, Lag-3, CD45, CD39, CD8⍺, (see details in Supplementary Table 14). Human CAR-T cells and TILs were stained with CD3, CD4, CD279, CD366(Tim-3), CD38, CD45, CD39, CD8⍺, CD233 (Lag-3), CD73, CD203a and TCF7 (Supplementary Table 14). Human patient derived cell lines were stained with anti-B7-H3 (Supplementary Table 14). Samples were acquired on a Cytek Northern Lights instrument and MA900 Sony Cell sorter and analysed with FlowJo software version 10.8.1.

### Analysis of mouse tumours and mouse T cells by flow cytometry

Single cell suspension from MDOTS processing (S3; see MDOTS generation section) was processed for flow cytometry). If frozen, single cell suspension was thawed into TexMACS media (Miltenyi Biotec, 130-097-196) and incubated at 37°C for 1 hour to allow cell recovery. The following fluorescently conjugated antibodies were used for mouse tumours and CD8^+^ TILs: CD3, CD4, CD8⍺, CD279, CD38, CD45, CD39,CD69, CD127, Lag-3, Tim-3 (see details in Supplementary Table 14). After washing, samples were acquired on a Cytek

Northern Lights instrument and analysed with FlowJo software.

### Measurement of mitochondrial membrane potential and mass

For flow cytometry analysis of mitochondrial membrane potential (MMP) and Mitochondrial mass T cells were stained with 10 nM MitoTracker Deep Red (ThermoFisher Scientific, M22426) and 10 nM MitoTracker Green (ThermoFisher Scientific, M7514), respectively for 30 min at 37 °C. Flow analysis was performed using Cytek Northern Lights instrument and mean fluorescence intensity (MFI) was analysed using FlowJo software version 10.8.1. For IncuCyte mitochondrial membrane potential analysis, plastic flat bottom 96 well plates were coated with 5 μg/mL human fibronectin in PBS (GIBCO, 14080-055) for 1h in 37 °C. Plates were washed 3 times in PBS and dried under sterile conditions. T cells were cultured in triplicates (30,000 per well) and allowed to adhere for 15 min. Mitochondrial membrane potential kit (Sartorius, 4775) was used for MMP measurements according to the manufacturer instructions. Cells were treated with 20 μM FCCP or 2.5 μg/mL Oligomycin, provided by the kit as controls. Cells and respective MMP measurements were acquired every 1 hour for 12h using Incucyte® Live-Cell Analysis System equipped with a Green/Orange/NIR, Metabolism and analysed using Incucyte® Cell-by-Cell Analysis Software Module (Cat. No. 9600-0031).

### Mitochondrial function analysis using Seahorse

Oxygen consumption rate (OCR) and extracellular acidification rate (ECAR) were determined using Seahorse Xfe96 analyzer (Agilent Technologies, Santa Clara, CA). Briefly, Seahorse XFe96/XF Pro FluxPak Mini culture plates (Agilent, 103793-100) were coated with 50 μg/mL Poly-Lysine (Thermo Scientific, A3890401) in water for 1h, washed with water and dried. Cartridge plate was hydrated with XF Calibrant (Agilent, 100840) for 4 hrs before experiment in a non-CO2 incubator. T cells (0.2 x 10^6^/well) were counted and plated in XF RPMI medium (pH 7.4, Agilent 103576) containing 10mM glucose (103577, Agilent), 2mM Glutamine (103579, Agilent) and 1mM pyruvate (103578, Agilent), followed by centrifugation to allow cell adhesion and monolayer formation. Cells were then placed in a non-CO2 incubator for 1hr. Mito Stress Test was performed using 1μM oligomycin (Sigma, 75351), 1.5μM FCCP (Sigma, C2920) and 100nM and 1μM, rotenone (sigma, R8875) and antimycin A (Sigma, A8674), respectively. OCR and ECAR rates were analysed using the Seahorse Wave Desktop software 2.6.

### Metabolite Profiling by mass spectrometry

*<u>Intracellular metabolomics:</u>* acutely or chronically stimulated CAR-T cells (see chronic stimulation section) or chronically stimulated +/- 3 days treatment of CD38 inhibitor 78c (6391, Toris) at 1uM (control DMSO) were analysed. At the day of the metabolomic analysis cells were cultured in fresh full RPMI media in equal concentrations for 2h. Four million cultured CAR-T cells (donor 1) or two million cultured CAR-T cells (donor 2), in three technical replicates were collected, centrifuged, washed with 0.9% NaCl, and resuspended in 300 μL extraction buffer (80% Methanol, 25 mM Ammonium Acetate and 2.5 mM Na-Ascorbate prepared in LC-MS water, supplemented with isotopically-labelled amino acid standards [Cambridge Isotope Laboratories, MSK-A2-1.2], aminopterin, and reduced glutathione standard [Cambridge Isotope Laboratories, CNLM-6245-10]). Samples were vortexed for 10 sec, then centrifuged for 10 minutes at 18,000g to pellet cell debris. The supernatant was dried on ice using a liquid nitrogen dryer. Dried samples were resuspended in 25 μL water and 2 μL was injected into a ZIC-pHILIC 150 x 2.1 mm (5 um particle size) column (EMD Millipore). operated on Vanquish™ Flex UHPLC Systems (Thermo Fisher Scientific, San Jose, CA, USA). Chromatographic separation was achieved using the following conditions: buffer A was acetonitrile; buffer B was 20 mM ammonium carbonate, 0.1% ammonium hydroxide in water; resulting pH is around 9 without pH adjustment. Gradient conditions we used were: 0–20 min: linear gradient from 20% to 80% B; 20–20.5 min: from 80% to 20% B; 20.5–28 min: hold at 20% B at 150 μL/min flow rate. The column oven and autosampler tray were held at 25 °C and 4 °C, respectively. MS data acquisition was performed using a QExactive benchtop orbitrap mass spectrometer equipped with an Ion Max source and a HESI II probe (Thermo Fisher Scientific, San Jose, CA, USA) and was performed in positive and negative ionisation mode in a range of m/z = 70–1000, with the resolution set at 70,000, the AGC target at 1 × 10^6^, and the maximum injection time (Max IT) at 20 msec. HESI settings were: Sheath gas flow rate: 35. Aux gas flow rate: 8. Sweep gas flow rate: 1. Spray voltage 3.5 (pos); 2.8 (neg). Capillary temperature: 320°C. S-lens RF level: 50. Aux gas heater temp: 350°C. *<u>Extracellular metabolomics:</u>* 100 μL of 3 days culture conditioned media of CAR-T cells cultures used for intracellular metabolomics (chronic/ chronic+CD38i) were collected and centrifuged at 1500 RPM for 5 min. Supernatant was collected and 30 μL of media was resuspended in 300 μL of lysis buffer (see intracellular section), followed by a similar downstream processing as the intracellular metabolomics.

### Data Analysis for metabolomics with TraceFinder

Targeted relative quantification of polar metabolites was standardly performed with TraceFinder 5.1 (Thermo Fisher Scientific, Waltham, MA, USA) using a 10 ppm mass tolerance and referencing an in-house library of chemical standards (230 compounds and 35 isotopically labelled internal controls). Pooled samples and fractional dilutions were prepared as quality controls and injected at the beginning and end of each run. In addition, pooled samples were interspersed throughout the run to control for technical drift in signal quality as well as to serve to assess the coefficient of variability (CV) for each metabolite. Data from TraceFinder was further consolidated and normalised with an in-house R script: (https://github.com/FrozenGas/KanarekLabTraceFinderRScripts/blob/main/MS_data_script_v2.4_20221018.R). Briefly, this script performs normalisation and quality control steps: 1) extracts and combines the peak areas from TraceFinder output .csvs; 2) calculates and normalises to an averaged factor from all mean-centred chromatographic peak areas of isotopically labelled amino acids internal standards within each sample; 3) filters out low-quality metabolites based on user inputted cut-offs calculated from pool reinjections and pool dilutions; 4) calculates and normalises for biological material amounts based on the total integrated peak area values of high-confidence metabolites. In this study, the linear correlation between the dilution factor and the peak area cut-offs are set to RSQ>0.95 and the coefficient of variation (CV) < 30%. Finally, Log2 fold change was calculated from the normalised peak areas of each metabolite relative to the first triplicate of the control and unpaired student *t*-test was performed, graphs were generated using GraphPad/Prism (v10.1.0) and unpaired *t*-test was applied. For PCA analysis and heat maps, data was normalised and Petro scaling was applied using MetaboAnalyst online platform. More information can be found in Supplementary Data Fig. 9-12.

### Data Analysis for metabolomics with CompoundDiscoverer

To expand our targeted analysis we performed relative quantification for polar metabolomics with CompoundDiscoverer (CD) 3.3 implementing additional in-house libraries (500 compounds). A general workflow was built to best suit our polar metabolomics LC-MS method. Positive and negative modes were analysed separately. Filtering steps were performed based on peak noise levels, ppm error, formula annotation, and the relative abundance of the integrated peak area in the true sample compared to blank injections, where features with +>3-fold higher in samples were retained. Within our HILIC chromatography, we rarely observe chromatographic shifts larger than 1 minute and retention times drifts larger than 40 seconds, thus these parameters were set as limits for retention time correction and matching to internal databases. Normalisation steps were performed based on internal standards and a strategy analogous to the one described above for TraceFinder. Post CD, positive and negative mode data were merged such that for metabolites detected in both, the mode at which higher values for total integrated peak area was retained. Downstream analysis was otherwise performed as above.

### NAD^+^ levels measurements

For NAD(H) measurements 1×10^5^ T cells were cultured in white 96 well plate and analysed using NAD/NADH-Glo™ luminescent assay (Promega, G9071), according to the manufacturer instructions. Luminescence was read every 10 min for 1h using Cytation 5 microplate reader and Gen5 software (BioTek). Relative NAD(H) levels were calculated as a fold change from control. Cells that were treated with 1uM 78c or 100uM NR were treated 1-3 days prior to the assay measurement.

### CoMut plots

Graphs were generated based on Crowdy et al.^59^, instructions can be found in Github https://github.com/vanallenlab/comut/blob/master/examples/documentation.ipynb. Python v3.9.13 was used to generate the plots.

### Statistical Methods, Data Analysis, and Software

All graphs with error bars report mean ± s.e.m. values except where indicated. Statistical tests, number of replicates, and independent experiments are listed in the text and figure legends. GraphPad/Prism (v10.1.0) was used for basic statistical analysis and plotting (http://www.graphpad.com). Schematics generated with BioRender (biorender.com) using a paid licence.

## Acknowledgements

This work was supported by a V Foundation Translational Research Grant (N.H., K.T.F, R.W.J.), NIH K08CA226391 (R.W.J.), Doris Duke Charitable Foundation Physician Scientist Fellowship (A.M.), NIH K12CA087723 (A.M), R01DE028172 (X.W.), R01CA226981 (X.W.), Department of Defense Idea Award W81XWH-20-PCRP-IDA (W81XWH2110433) (X.W.), N.K is a Pew Scholar, NIH K08CA234458, Doris Duke Clinical Scientist Training Grant (D.L). Additional support provided by the Termeer Early Career Fellowship in Systems Pharmacology (R.W.J.), and a generous gift from Robert and Marie McInnes. We acknowledge funding provided by Massachusetts Life Sciences Center Research Infrastructure Program in support of the Mass General Cancer Center Tumour Cartography Center, and the Dr. Miriam and Sheldon G. Adelson Medical Research Foundation. The funding bodies had no role in the design of the study, and collection, analysis, and interpretation of the data, or in writing the manuscript. The authors thank all members of the Hacohen and Jenkins laboratories at MGH and the Broad Institute. Graphics in Fig. 2g, 3a, and 4q were created with Biorender.com using a paid licence.

## Author Contributions

*Conception and experimental design*: O.Y.R., A.M.C., M.S.F., N.H., J.M.S., L-C.C, N.S., R.S., and R.W.J. Methodology and data acquisition: O.Y.R., A.M.C, B.P., J.Y., N.M., L.C.C, N.S., E.E., M.M., S.A.E., N.N., M.E.W., R.S., and M.S.F. *Analysis and interpretation of data*: O.Y.R., O.S., S.A., J.P., L.C., A.M., S.J.W., L.B., K.Y., J.Y., J.M.S., L.C.C, N.S., E.E., M.M., N.N., M.E.W., M.H.S, N.K., M.S.F, N.H., A.M.C., and R.W.J. *Administrative, technical, or material support*: G.C., A.T.M, P.T, T.S., A.L., H.X., W.A.M., M.Q.R., J.F., C.P., F.C., X.W., C.R.F., D.L., K.T.F., D.R.S., U.M.S., R.T.M., D.P.L., R.J.S., Y.S., M.K., and G.M.B. *Manuscript writing and revision*: O.Y.R., M.S.F., N.H, O.S, B.P., N.K., S.A, J.P, S.J.W, A.M., J.Y. and R.W.J.

## Competing Interests

R.W.J. is a member of the advisory board for and has a financial interest in Xsphera Biosciences Inc., a company focused on using ex vivo profiling technology to deliver functional, precision immune-oncology solutions for patients, providers, and drug development companies. R.W.J. has received honoraria from Incyte (invited speaker), G1 Therapeutics (advisory board), Bioxcel Therapeutics (invited speaker). R.W.J. has ownership interest in U.S. patents US20200399573A9 and US20210363595A1. R.W.J.’s interests were reviewed and are managed by Massachusetts General Hospital and Mass General Brigham in accordance with their conflict-of-interest policies. A.M has served a consultant/advisory role for Third Rock Ventures, Asher Biotherapeutics, Abata Therapeutics, ManaT Bio, Flare Therapeutics, venBio Partners, BioNTech, Rheos Medicines and Checkmate Pharmaceuticals, is currently a part-time Entrepreneur in Residence at Third Rock Ventures, is an equity holder in ManaT Bio, Asher Biotherapeutics and Abata Therapeutics, and has received research funding support from Bristol-Myers Squibb. A.M.’s interests were reviewed and are managed by Massachusetts General Hospital and Mass General Brigham in accordance with their conflict-of-interest policies. J.M.S., L-C.C, N.S, M.M., N.N., R.S. are current employees with Teiko.bio and own stock. E.E. and S.A.E. were employed with Teiko.bio in the past 2 years and own stock. R.S. and M.H.S. are Teiko.bio’s co-founders and R.S. serves on the company board. M.H.S. serves as an advisor for Teiko.bio and owns stock. M.E.W. worked as a contractor for Teiko.bio during this project. M.H.S. has received a speaking honorarium from Standard BioTools and Kumquat Bio, has been a paid consultant for Five Prime, Ono, January, Earli, Astellas, and Indaptus, and has received research funding from Roche/Genentech, Pfizer, Valitor, and Bristol Myers Squibb.X.W. and C.R.F. report a patent on the B7-H3 CAR T cells (US10519214B2). K.T.F. serves on the Board of Directors of Clovis Oncology, Strata Oncology, Kinnate, and Scorpion Therapeutics; Scientific Advisory Boards of PIC Therapeutics, Apricity, C-Reveal, Tvardi, ALX Oncology, xCures, Monopteros, Vibliome, Karkinos, Soley Therapeutics, Alterome, Immagene, and intrECate; consultant to Nextech, Takeda, Novartis, Transcode Therapeutics, and Roche/Genentech. R.T.M. consults for Bristol Myers Squibb. G.M.B. has sponsored research agreements through her institution with: Olink Proteomics, Teiko Bio, InterVenn Biosciences, and Palleon Pharmaceuticals. She has served on advisory boards for: Iovance, Merck, Nektar Therapeutics, Novartis, and Ankyra Therapeutics. She consults for: Merck, InterVenn Biosciences, Iovance, and Ankyra Therapeutics. She holds equity in Ankyra Therapeutics. M.S.F received funding from Calico Life Sciences, Bristol-Myers Squibb, Istari Oncology and served as a consultant for Galvanize Therapeutics. N.H. holds equity in BioNTech and is an advisor for Related Sciences/Danger Bio, Repertoire Immune Medicines and CytoReason, and receives research funding from Calico Life Sciences and Bristol-Myers Squibb. D.L serves on the scientific advisory board for Oncovalent Therapeutics, and received honorariums from Genentech.

**Correspondence and requests for materials** should be addressed to R.W.J.

## Dedication

The authors would like to dedicate this work to the memory of the late Soldano Ferrone, MD PhD.

## Extended Data Figure Legends

**Extended Figure 1.**
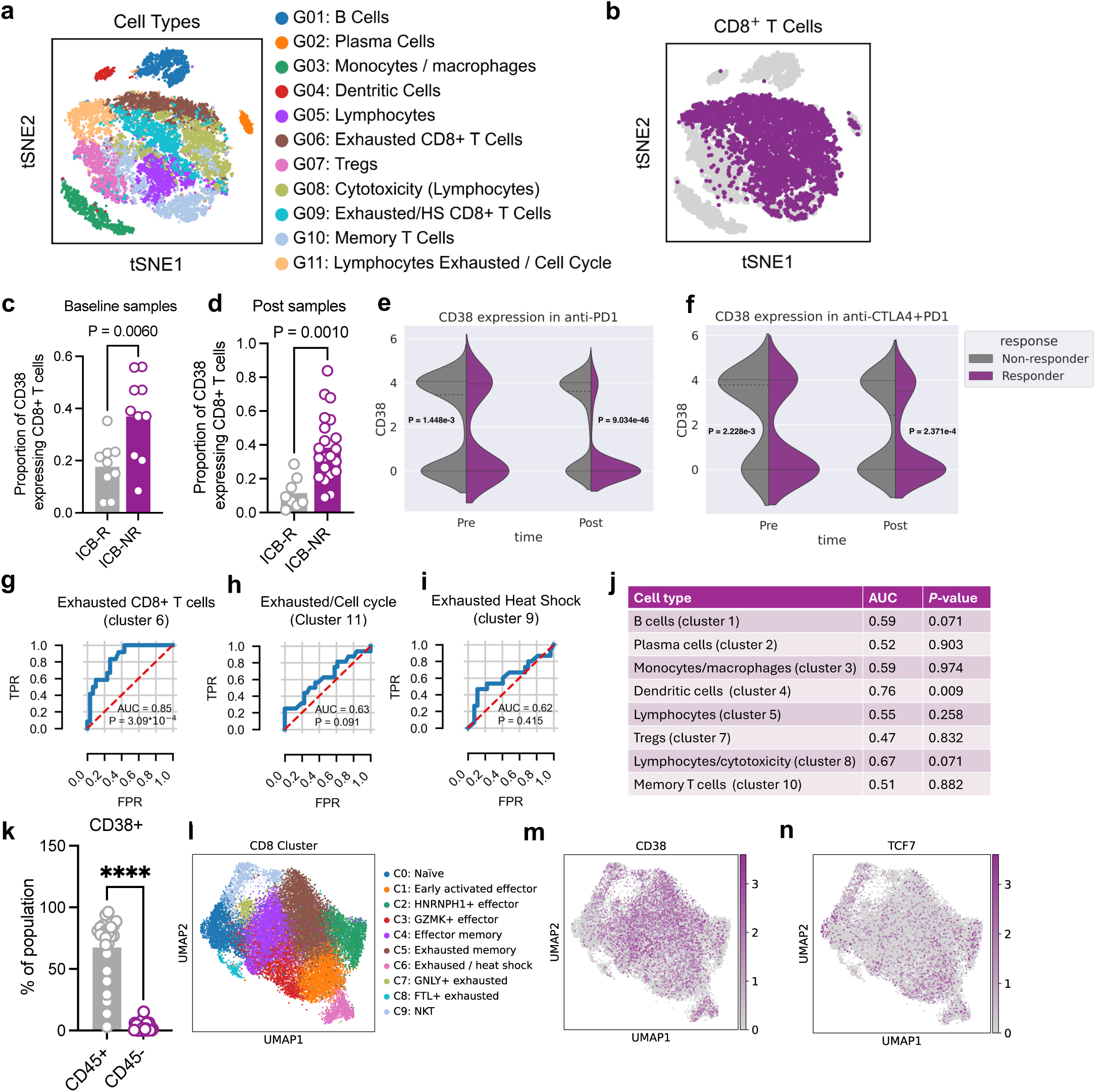
Supporting data that CD38^+^ CD8^+^ T cells are associated with ICB resistance. **a-d,** scRNAseq of tumour-infiltrating leukocytes (CD45^+^) from melanoma patients treated with ICB^10^ **(a)** t-SNE plot of CD45^+^ (n=16,291) with 11 distinct clusters; **(b)** feature plot demonstrating *CD8a* expression; **(c)** proportion of *CD38* expressing CD8^+^ T cells from pre-ICB tumours; (ICB-R, *n*=9; ICB-NR, *n*=10) and **(d)** post ICB tumours (ICB-R, *n*=8; ICB-NR, *n*=21; Means (bars) and individual values (open circles) are shown; 2-sided unpaired *t*-test. **e-f,** proportion of *CD38* expressing CD8 T cells in ICB-R and ICB-NR, separated by ICB type (anti-PD-1 or anti-PD-1 plus anti-CTLA-4), (ICB-R, *n*=17; ICB-NR, *n*=31) nonparametric two sample Kolmogorov-Smirnov test. **g-i,** Receiver Operating Characteristic (ROC) curves demonstrating the predictive power of CD38^+^CD45^+^ cells for ICB resistance across different immune clusters in Extended Data Fig. 1a (clusters 6,9,11). FPR = false positive rate. TPR = true positive rate. **j**, summary of Receiver Operating Characteristic (ROC) curves of the predictive power of *CD38* expression across different CD45^+^ clusters for ICB resistance. **k**, flow cytometry immunophenotyping of CD38 surface staining in CD45^+^ and CD45^-^ cells from human melanoma tumours (*n=*18). Means (bars) and individual values (open circles) are shown; 2-sided unpaired *t*-test. **l-n,** scRNAseq analysis of CD8 T cells from melanoma validation cohort **(l)** UMAP of CD8 clusters (n=22,008) with 9 distinct clusters identified; **(m)** *CD38* expression **(n)** *TCF7* expression. \*\*\*\**P* < 0.0001.

**Extended Figure 2.**
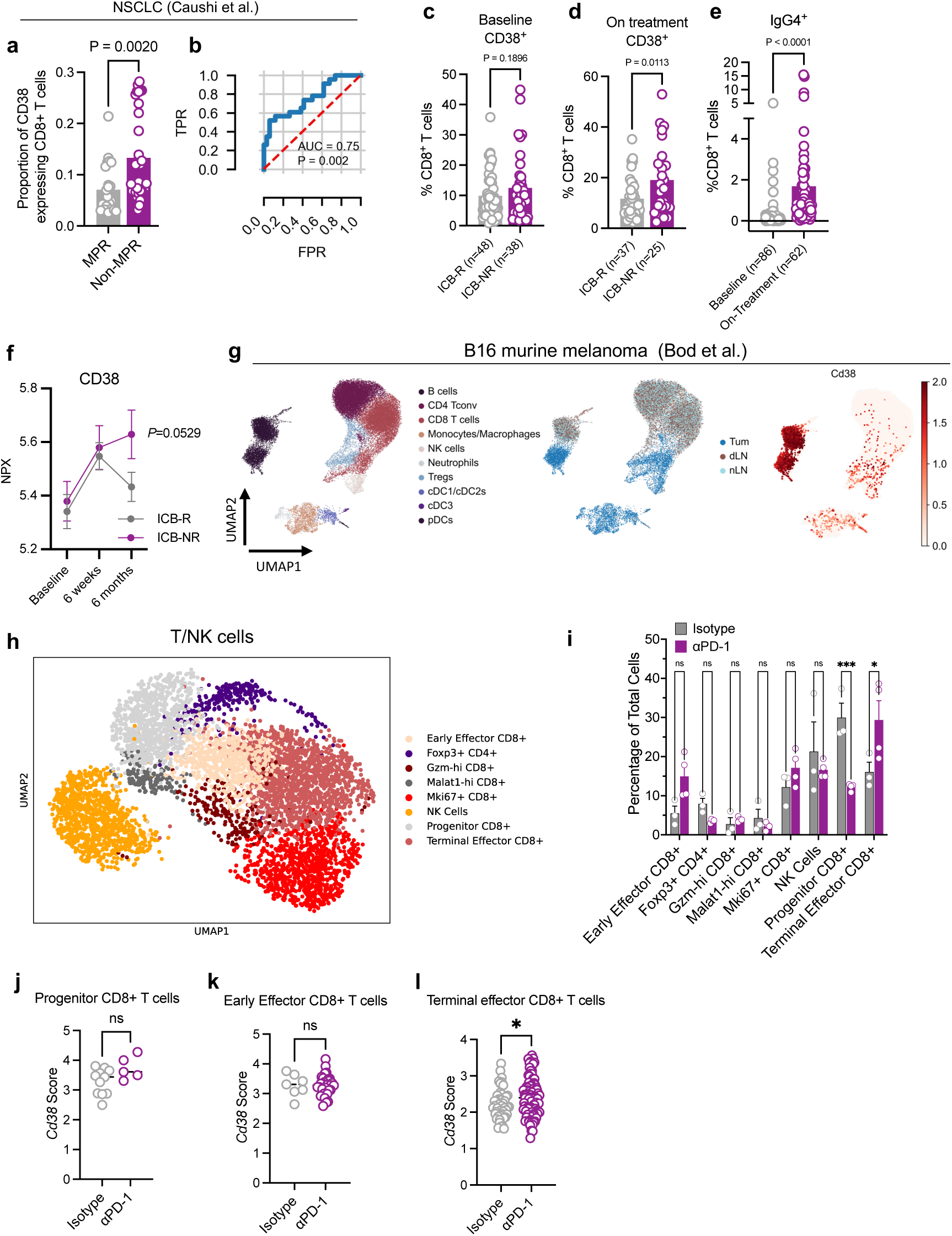
Supporting data that CD38^+^ CD8^+^ T cells are increased during tumour progression. **a,** proportion of *CD38* expressing CD8^+^ T cells from **(a)** NSCLC^18^ (MPR-major pathological response ICB-R *n*=23; non-MPR, ICB-NR, *n*=34) **b,** Receiver Operating Characteristic (ROC) curve demonstrating the predictive power of CD38^+^CD8^+^ T cells for lack of ICB treatment benefit in NSCLC; FPR = false positive rate. TPR = true positive rate. **c-d** CyTOF analysis of peripheral blood examining CD8^+^CD38^+^ T cells in melanoma patients’ blood before ICB treatment **(c)** and after **(d)**. **e**, CyTOF analysis of peripheral blood examining CD8^+^CD38^+^IgG4^+^. **f,** Olink analysis of CD38 protein levels in blood plasma of ICB-R (*n*=64) and ICB-NR (*n*=53) patients before and following 6 weeks and 6 months after ICB treatment. 2-way ANOVA. **g,** scRNAseq of CD45^+^ cells from B16 tumours showing UMAP with 10 distinct clusters, their distribution of location (Tum), tumour draining lymph nodes (dLN), and normal lymph nodes (nLN) and the expression of *CD38* in those clusters. **h**, scRNAseq of T/NK tumour-infiltrating leukocytes from B16-ova tumours from Control (Vehicle/IgG, n=3) and αPD-1 (Vehicle/anti-PD-1, n=4). **i-l,** CD3^+^ TILs in Control (Vehicle/IgG, n=3) and αPD-1 (Vehicle/anti-PD-1, n=4) B16-ova treated tumours. **(i)** proportion of indicated immune populations upon PD-1 blockade (proportion from total cells; 2-way ANOVA with Sidak correction for multiple comparisons.). **j-l,** proportion of *Cd38* expressing CD3^+^ TILs of indicated types; unpaired *t*-test. \**P* <0.05, \*\*\**P* <0.001.

**Extended Figure 3.**
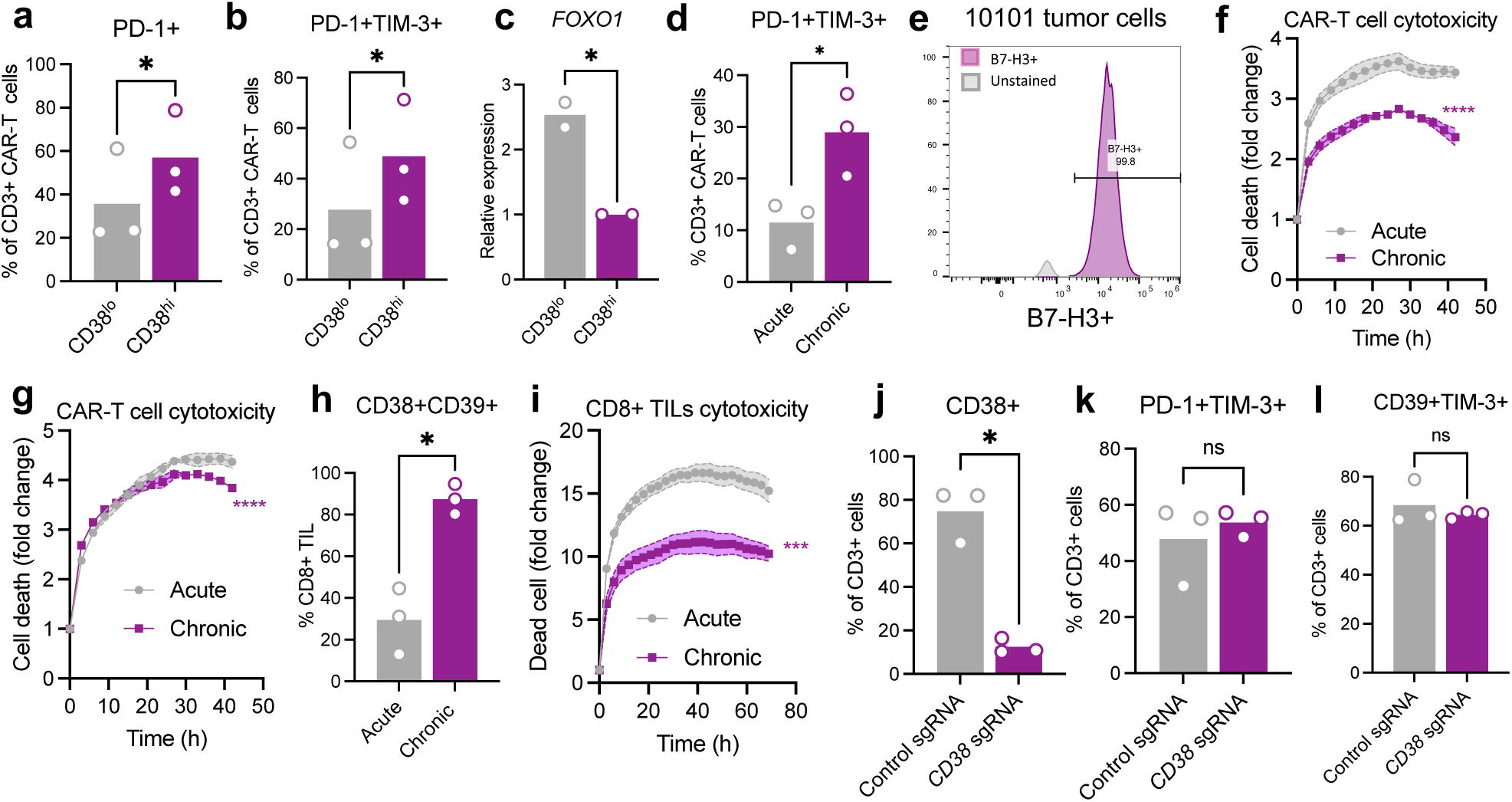
Supporting data that CD38^hi^CD8^+^ T cells are dysfunctional. **a-c,** analysis of sorted CD38^hi^ and CD38^lo^ B7-H3.CAR-T for **(a)** surface staining for PD-1^+^, **(b)** surface staining of PD-1^+^TIM-3^+^ and **(c)** gene expression of *FOXO1*. Means (bars) and individual values (open circles) are shown (*n*=3 (a-b); *n*=2 (c); 2-sided unpaired *t*-test). **d,** analysis of surface staining for PD-1^+^TIM-3^+^ in B7-H3.CAR-T cells following acute and chronic T cell stimulation. Means (bars) and individual values (open circles) are shown (n=3; 2-sided paired *t*-test). **e,** Analysis of surface staining for B7-H3 antigen in 10101 melanoma tumour cells **f-g,** cytotoxicity assay of acute and chronically stimulated B7-H3.CAR-T cells towards 10170 **(f)** and 10101 **(g)** patient derived melanoma cell lines. Two other repeats for Fig. 2k. Means +/- s.e.m. (shaded area) are shown (n=3 biological replicates; 2-way ANOVA with Sidak correction for multiple comparisons). **h,** surface staining for CD38^+^CD39^+^ in CD8^+^ TILs from melanoma patients, following acute and chronic T cell stimulation. Means (bars) and individual values (open circles) are shown (n=3; 2-sided paired *t*-test) **i,** cytotoxicity of 10170 CD8^+^ TILs following acute and chronic T cell stimulation, towards 10170 patient-derived cell line (n=2 biological replicates; 2-way ANOVA with Sidak correction for multiple comparisons). **j-l,** analysis of surface staining of **(j)** CD38^+^, **(k)** PD-1^+^TIM-3^+^, **(l)** CD39^+^TIM-3^+^ in chronically stimulated control sgRNA and *CD38* sgRNA B7-H3.CAR-T cells. Means (bars) and individual values (open circles) are shown (n=3; 2-sided paired *t*-test). \**P* <0.05, \*\*\**P* < 0.001, \*\*\*\**P* < 0.0001, *ns*, not significant.

**Extended Figure 4.**
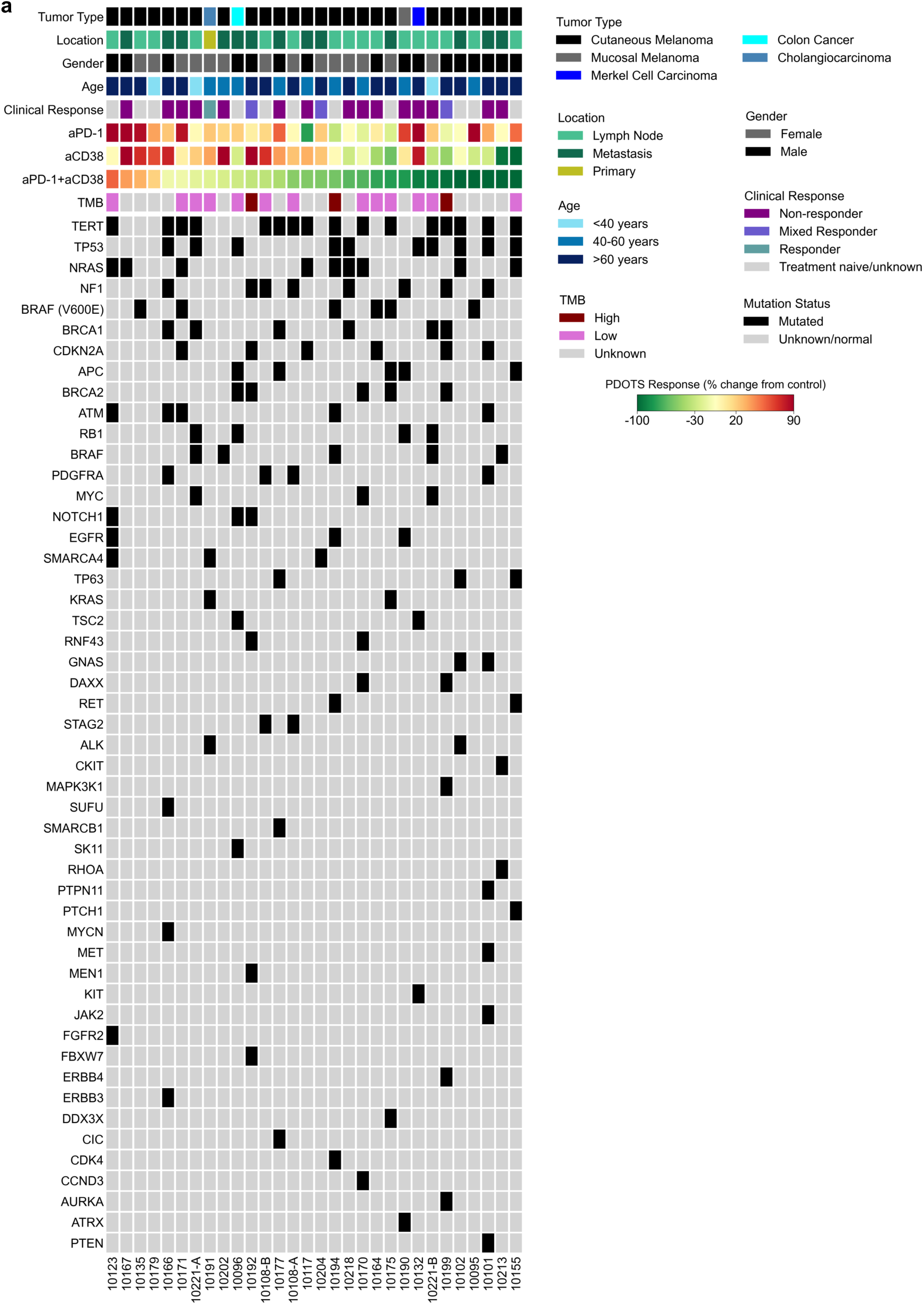
Supporting data that CD38 blockade overcomes ICB resistance. **a**, Tumour type, tissue source (location), gender, age, clinical response, PDOTS response data, tumour mutational burden (TMB), and associated tumour mutational profile for specimens used for PDOTS profiling (*n*=30; samples ordered by response to dual PD-1/CD38 blockade). PDOTS response parameters defined as previously described^21^: responder (reduction >30% compared to control), partial responder (<30% reduction and <20% growth compared to control), and non-responder (>20% growth compared to control).

**Extended Figure 5.**
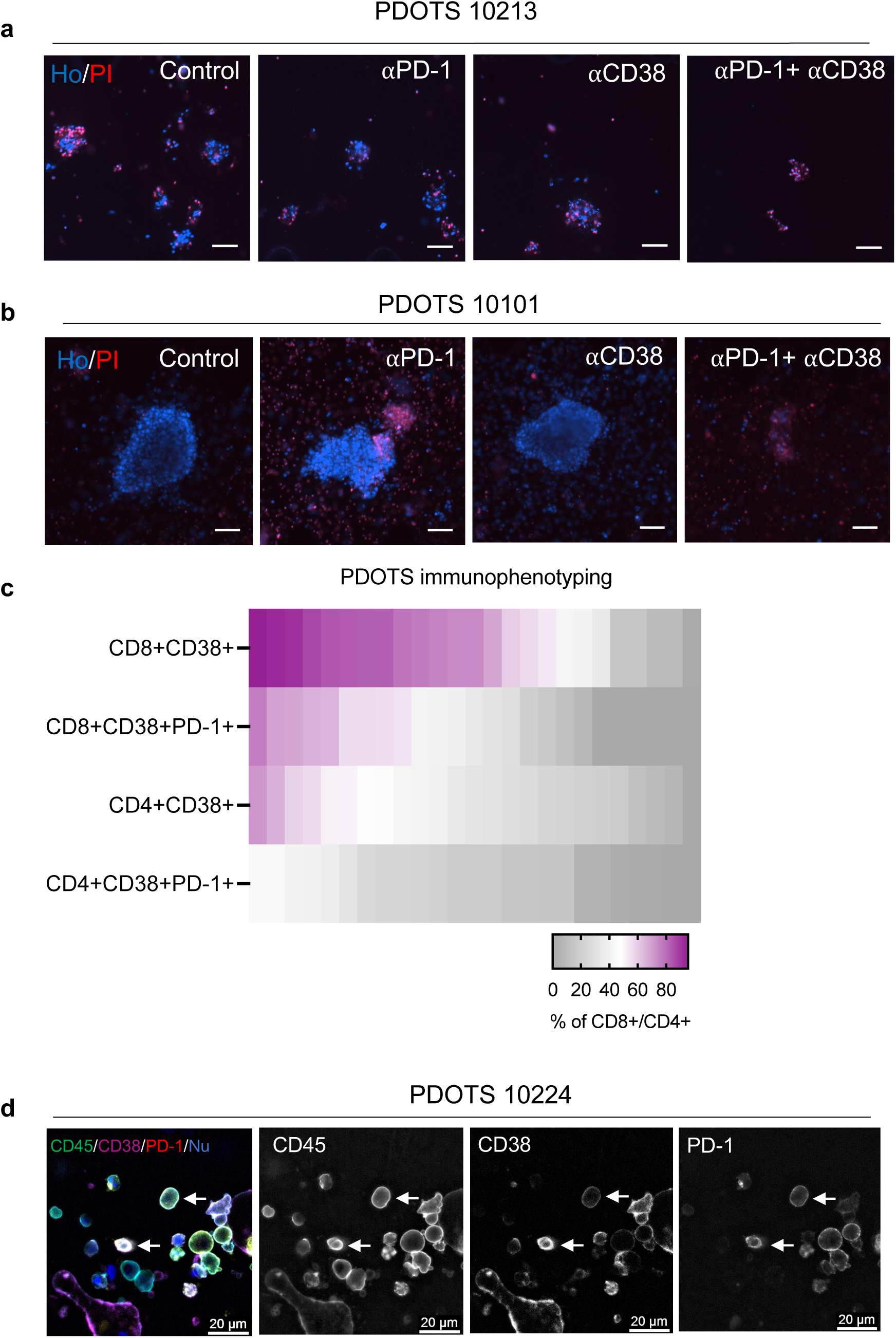
Supporting data that CD38 blockade overcomes ICB resistance in human melanoma. **a-b**, representative images of 10213 ICB-resistant mucosal melanoma PDOTS **(a)** and 10101 ICB-resistant cutaneous melanoma PDOTS **(b)** treated with indicated treatments. Ho-Hoechst (blue), PI -propidium iodide, dead cells (red). Scale bars=100μm. **c,** flow cytometry analysis of indicated populations from PDOTS single cell suspension before culturing (*n*=25). **d,** immunofluorescence imaging of cutaneous melanoma PDOTS with indicated markers. Nu-nucleus, blue. White arrows denote cells with marker co-expression.

**Extended Figure 6.**
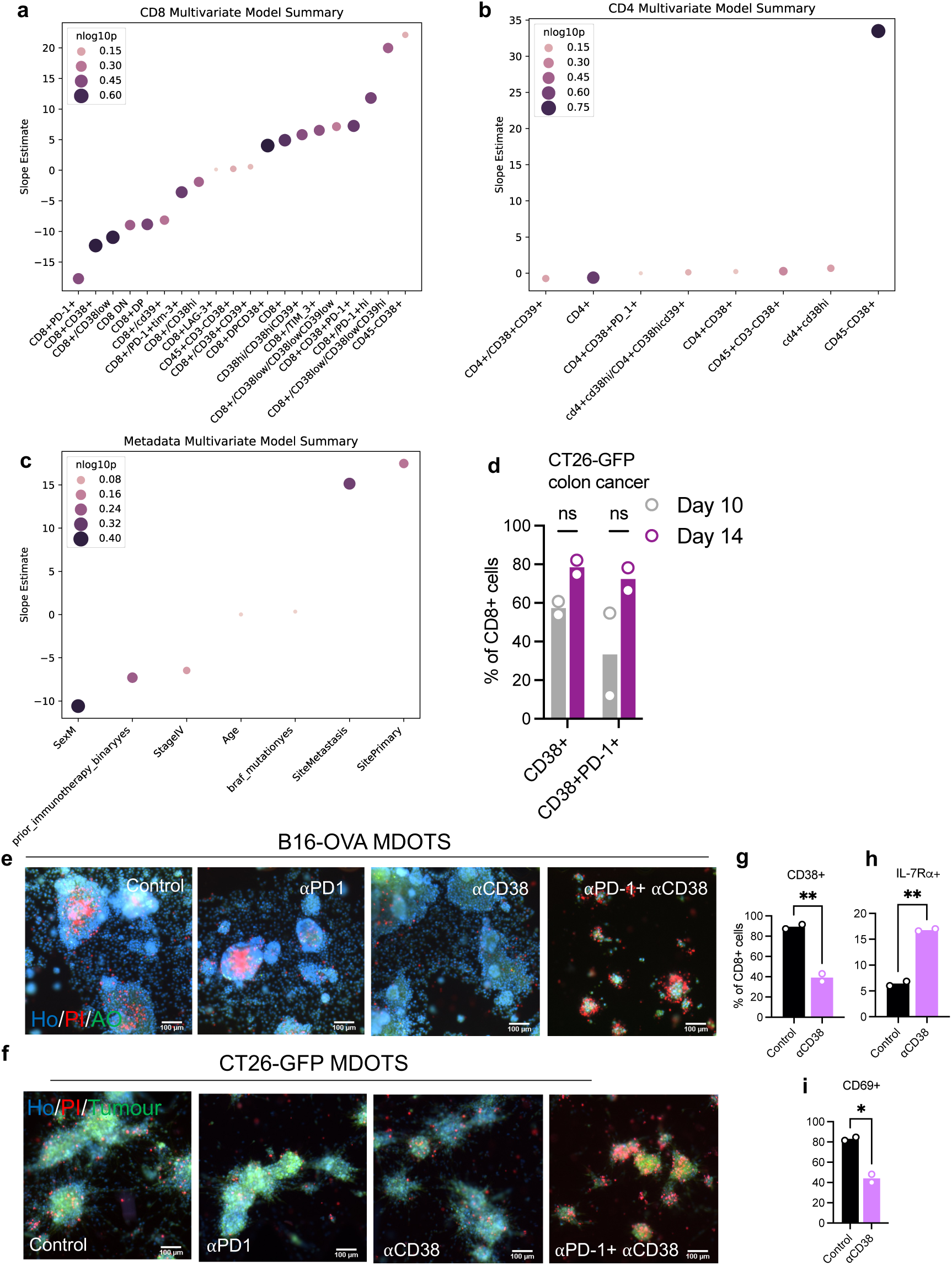
Supporting data that CD38 blockade overcomes ICB resistance. **a-c,** Multiple linear regression summaries for the association of immune and clinical features of melanoma patient derived tumours used for PDOTS with *ex vivo* response to PD-1/CD38 dual blockade (*n*=30 for clinical features and *n*=22 for immune features) (Fig. 3b-c). **(a)** flow cytometry measures CD8^+^ proportions; **(b)** CD4^+^ proportions; **(c)** clinical demographic features. Points are coloured by -log10(p-value). **d,** flow cytometry analysis of indicated populations in CD8^+^ T cells TILs from CT26-GFP tumours at day 10 and day 14**. e-f,** representative images of B16-ova murine melanoma **(e)** and CT26-GFP murine colon cancer **(f)** treated with indicated treatments. Ho-Hoechst (blue), PI-PI -propidium iodide, dead cells (red), AO-acridine orange, live cells (green). **g-i**, cell surface staining of **(g)** CD38^+^ **(h)** IL-7Ra^+^ **(i)** CD69^+^ on CT26-GFP murine tumour derived CD8^+^ TILs treated *in vitro* with CD38 blocking antibody or control IgG. Means (bars) and individual values (open circles) are shown (*n*=2; 2-sided unpaired *t*-test). \**P* <0.05, \*\**P* < 0.01.

**Extended Figure 7.**
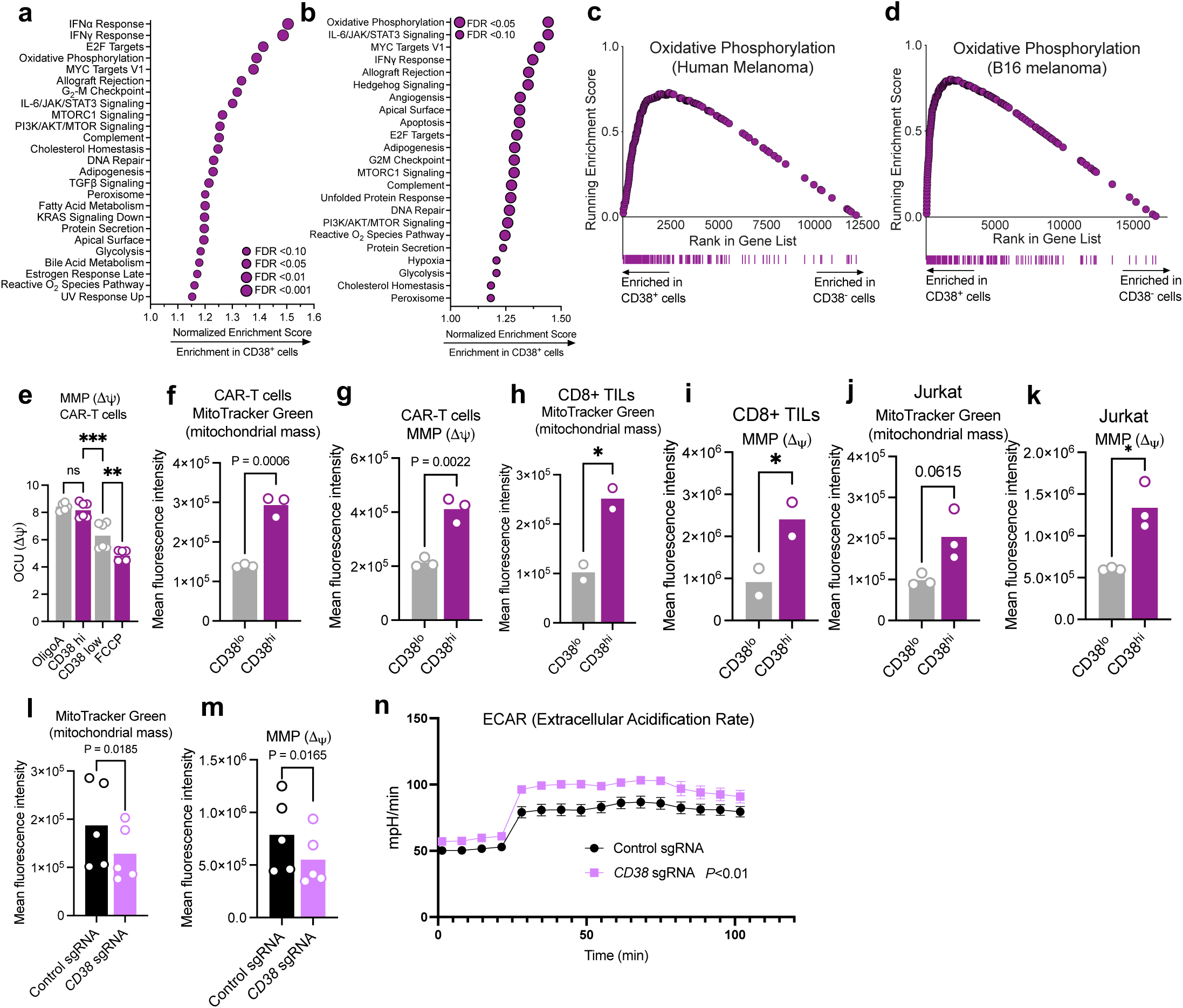
Supporting data that CD38 is associated with mitochondrial dysfunction. **a-b,** Gene set enrichment analysis (GSEA) of **(a)** CD8^+^ TILs from human melanoma^10^ and **(b)** CD3^+^ TILs from B16-ova murine melanoma^21^**. c-d,** mountain plots showing enrichment for OXPHOS gene set based on *CD38* expression in (**c**) CD8^+^ TILs human melanoma^10^ and (**d**) CD3^+^ TILs from B16-ova murine melanoma^21^. **e,** Incucyte mitochondrial membrane potential (MMP) measurements of sorted CD38^hi^ and CD38^lo^ B7-H3.CAR-T cells. Means (bars) and individual values (open circles) are shown (n=3; 3 biological replicates; 1-way ANOVA). **f-k,** Flow cytometry analysis of mitochondrial mass (**f,h,j**) and MMP (**g,i,k**) in CD38^hi^ and CD38^lo^, chronically stimulated B7-H3.CAR-T cells (**f,g**), CD8^+^ TILs (**h,i**) and JURKAT cells (**j,k**). Means (bars) and individual values (open circles) are shown (n=3; 2-sided paired *t*-test). **l,m,** Flow cytometry analysis of mitochondrial mass (**l**) and MMP (**m**) of chronically stimulated control and *CD38* sgRNA B7-H3.CAR-T cells (*n*=3; Means (bars) and individual values (open circles) are shown; 2-sided unpaired *t*-test). **n,** Extracellular acidification rate (ECAR) under basal condition in response to addition of oligomycin, carbonyl cyanide *p*-trifluoromethoxyphenylhydrazone (FCCP), and rotenone/antimycin A of B7-H3.CAR-T cells form indicated groups. (n=5, 2 biological replicates, 2-way ANOVA with Sidak correction for multiple comparisons). \**P* <0.05, \*\**P* < 0.01, \*\*\**P* < 0.001.

**Extended Figure 8.**
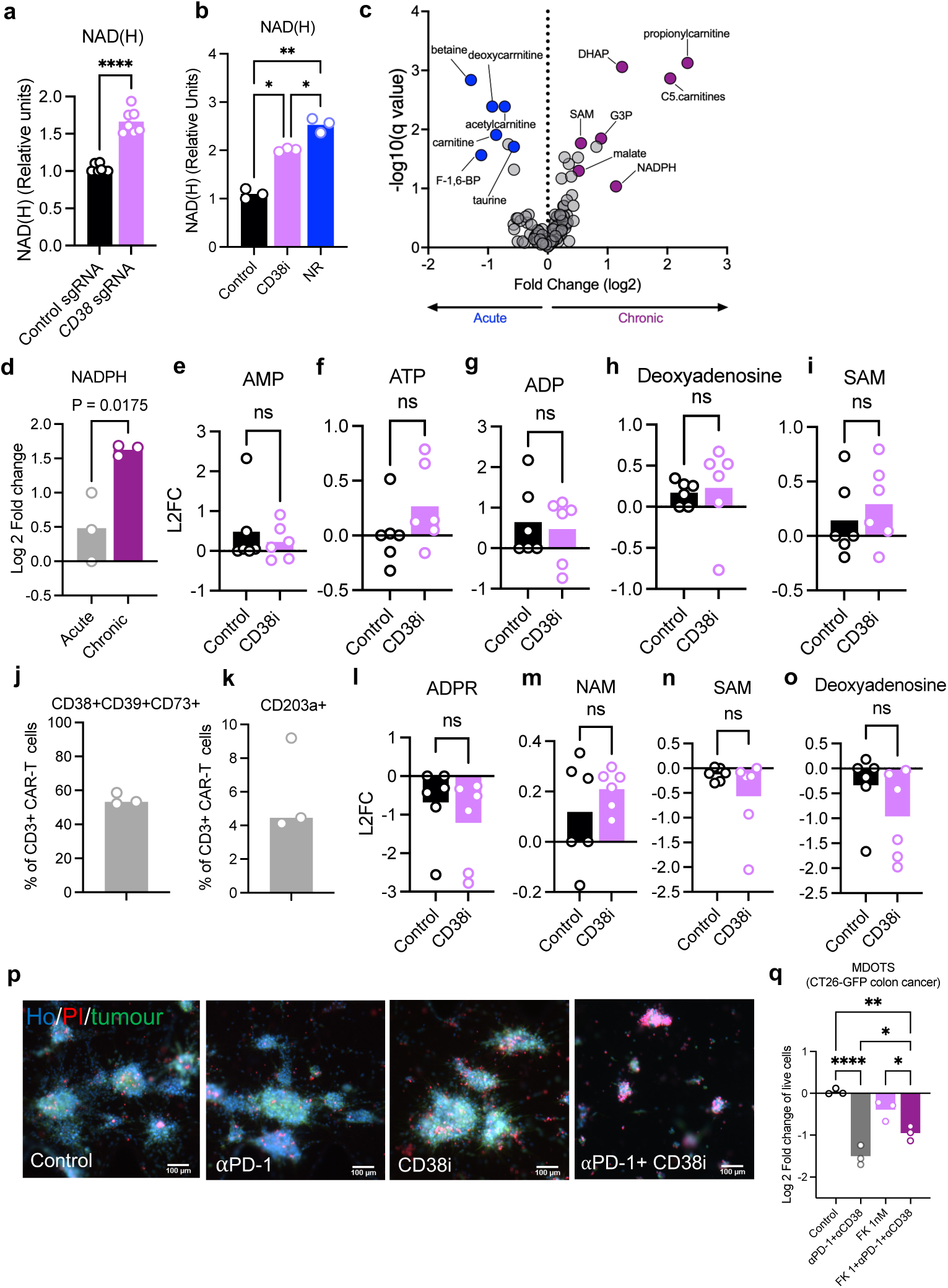
Supporting data that CD38 inhibition restores cellular NAD^+^ levels. **a-b,** Analysis of relative NAD(H) in (**a**) control and *CD38* sgRNA B7-H3.CAR-T cells (n=3 biological repeats, 3 independent experiments; Means (bars) and individual values (open circles) are shown; 2-sided unpaired *t*-test) and (**b**) chronically stimulated B7-H3.CAR-T cells with indicated treatments (n=3 biological repeats; Means (bars) and individual values (open circles) are shown; 1-way ANOVA with Tukey correction for multiple comparisons). **c,** volcano plot of intracellular metabolomics of acute vs chronic B7-H3.CAR-T cells (n=3 biological repeats). **d,** Relative NADPH levels in B7-H3.CAR-T cells in indicated groups determined by LC-MS. Means (bars) and individual values (open circles) are shown (n=3 biological replicates, 2-sided unpaired *t*-test). **e-i,** Relative levels of intracellular **(e)** AMP (Adenosine mono-phosphate), **(f)** ATP (Adenosine triphosphate), **(g)** ADP(Adenosine diphosphate), **(h)** deoxyadenosine and **(i)** SAM (S-Adenosylmethionine) in chronically stimulated CAR-T cells +/- CD38i (n=6; 2 independent CAR-T donors; Means (bars) and individual values (open circles) are shown; 2-sided unpaired *t*-test). **j-k,** flow cytometric analysis of the indicated surface markers in chronically stimulated B7-H3.CAR-T cells (n=3 biological repeats; Means (bars) and individual values (open circles) are shown). **l-o,** Relative levels of extracellular **(l)** ADPR (Adenosine diphosphate ribose), **(m)** NAM (nicotinamide), **(n)** SAM, **(o)** Deoxyadenosine in chronically stimulated CAR-T cells +/- CD38i (n=6 biological replicates; 2 independent CAR-T donors; 2-sided unpaired *t*-test). **p,** representative images of CT26-GFP murine colon cancer treated with indicated treatments. Ho-Hoechst (blue), PI- PI -propidium iodide, dead cells (red), GFP- tumour cells. **q,** Viability assessment of CT26-GFP MDOTS (n=3 biological replicates, one-way ANOVA with Tukey correction for multiple comparisons), with indicated treatments. \**P* < 0.05, \*\**P* < 0.01, \*\*\**P* < 0.001, *ns*, not significant.

## Supplemental Information Guide

**Supplemental Table 1-** Melanoma validation cohort. Fig. 1h.

**Supplemental Table 2-** Melanoma patients blood cohort Fig. 1i-j, Extended data Fig. 2c-e.

**Supplemental Table 3-** Differential gene expression analysis of CD38^+^ vs CD38-CD8^+^ T cells in human melanoma. Fig. 2c.

**Supplemental Table 4-** Differential gene expression analysis of CD38^+^ vs CD38-CD3^+^ TILs from B16-OVA murine melanoma. Fig. 2d.

**Supplemental Table 5-** PDOTS cohort demographics (pembrolizumab, daratumumab). Fig. 3b-c, Extended Data Fig. 4a.

**Supplemental Table 6 -** Gene Set Enrichment Analysis of CD38^+^/CD38^-^ CD8+ TILs in human melanoma Fig. 4a, Extended data Fig. 7a,c.

**Supplemental Table 7 -** Gene Set Enrichment Analysis of CD38^+^/CD38-CD3^+^ TILs in B16 murine melanoma Fig. 4a, Extended data Fig. 7b,d

**Supplemental Table 8-** Intracellular metabolomics of acute and chronically stimulated B7.H3 CAR-T cells. Extended data Fig. 8c-d.

**Supplemental Table 9-** Intracellular metabolomics of chronically stimulated +/- CD38i B7.H3 CAR-T cells. 4d-h, Extended data Fig. 8e-i.

**Supplemental Table 10-** Extracellular metabolomics of chronically stimulated +/- CD38i B7.H3 CAR-T cells. Extended data Fig. 8l-o.

**Supplemental Table 11-** PDOTS cohort demographics (pembrolizumab, CD38i, NR). Fig. 4o-p.

**Supplemental Table 12-** CyTOF analysis antibodies list. Fig. 1i-j, Extended data Fig. 2c-e.

**Supplemental Table 13-** CD38 CRISPR guides RNA sequences.

**Supplemental Table 14-** flow cytometry and immunofluorescence antibodies list.

**Supplemental Table 15-** primers sequences for qPCR.

## Supplemental Figures -

**Supplementary Data Fig. 1- a,** single cell RNAseq tSNE plots of all CD45^+^ cells and mapping of ICB response. **c-j,** (ROC) analysis showing the ICB resistance predictive power of CD38^+^CD8^+^ T cells from different clusters in human scRNA seq. Supporting Extended Data Fig. 1j.

**Supplementary Data Fig. 2- a-d,** Supporting Extended Data Fig. 1k. (a-b) single cell RNAseq tSNE plots of melanoma tumour cells clusters and feature plot demonstrating *CD38* expression in melanoma cells^60,61^. **c,** *CD38* expression in tumours from TCGA dataset versus normal tissue. Melanoma (SKCM) is highlighted in purple. **d,** *CD38* expression in melanoma cell lines from the CCLE database, clustered against proliferative melanoma (*MITF* high) versus neural crest like melanoma (*AXL* high). **e,** T cell markers that were used for the classification of T cell clusters (Extended Data Fig. 1l).

**Supplementary Data Fig. 3- a,** gating strategy for patient blood CD8^+^ T cells CyTOF analysis. Supporting Fig. 1i-j, Extended Data Fig. 2c-e.

**Supplementary Data Fig. 4- a,** gating strategy for B7.H3 CAR-T cells. **b,** flow analysis of CD38 CRISPR edited B7.H3 CAR-T cells. **c-d,** B7.H3 CAR-T cells CD38 hi/low sorting strategy.

**Supplementary Data Fig. 5-** PDOTS CD4^+^ and CD8^+^ gating strategy. Supporting Extended Data Fig. 5c.

**Supplementary Data Fig. 6-** Mouse tumours gating strategy. Supporting Extended Data Fig. 6d.

**Supplementary Data Fig. 7-** Mouse CD8^+^ isolated TILs gating strategy. Supporting Extended Data Fig. 6g-i.

**Supplementary Data Fig. 8-** Mitochondrial stain gating strategy. Supporting Extended Data Fig. 7f-m.

**Supplementary Data Fig. 9-** Intracellular metabolomics of acute versus chronically stimulated CAR-T cells. Supporting Extended Data Fig. 8c-d.

**Supplementary Data Fig. 10-12-** Intracellular and extracellular metabolomics of chronically stimulated CAR-T cells +/- CD38i. Supporting Fig. 4d-h, Extended Data Fig. 8e-i. 8l-o.

## Notes

### Summary of Updates

Added a clarification for one of the references in the introduction and Fig.1 main text.

