## Supplementary Material for "Disrupting CD38-driven T cell dysfunction restores sensitivity to cancer immunotherapy"

Supplementary Data Figure 1

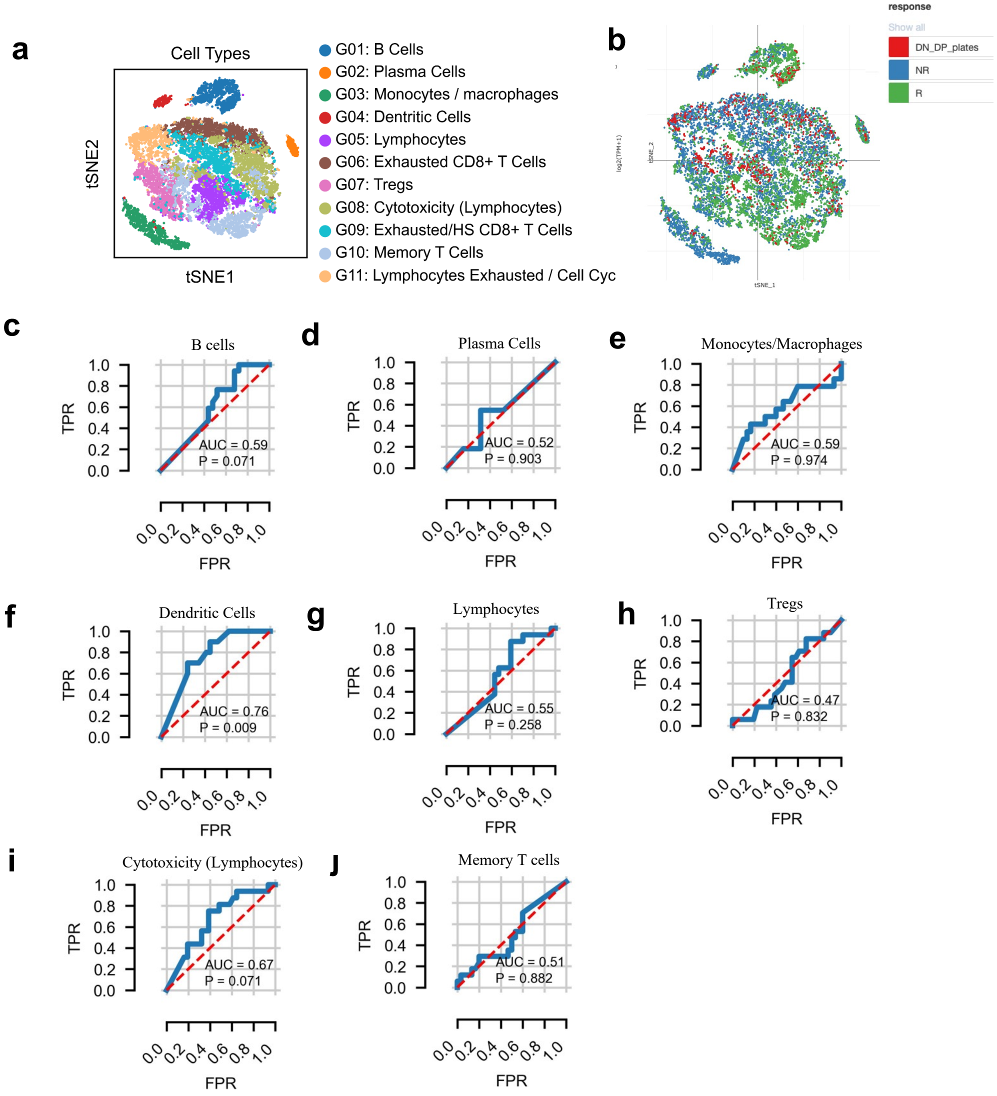


**Supplementary Data Fig. 1- a,** single cell RNAseq tSNE plots of all CD45+ cells and mapping of ICB response. **c-j,** (ROC) analysis showing the ICB resistance predictive power of $CD38^{+}CD8^{+}$ T cells from different clusters in human scRNA seq. Supporting Extended Data Fig. 1j.

Supplementary Data Figure 2

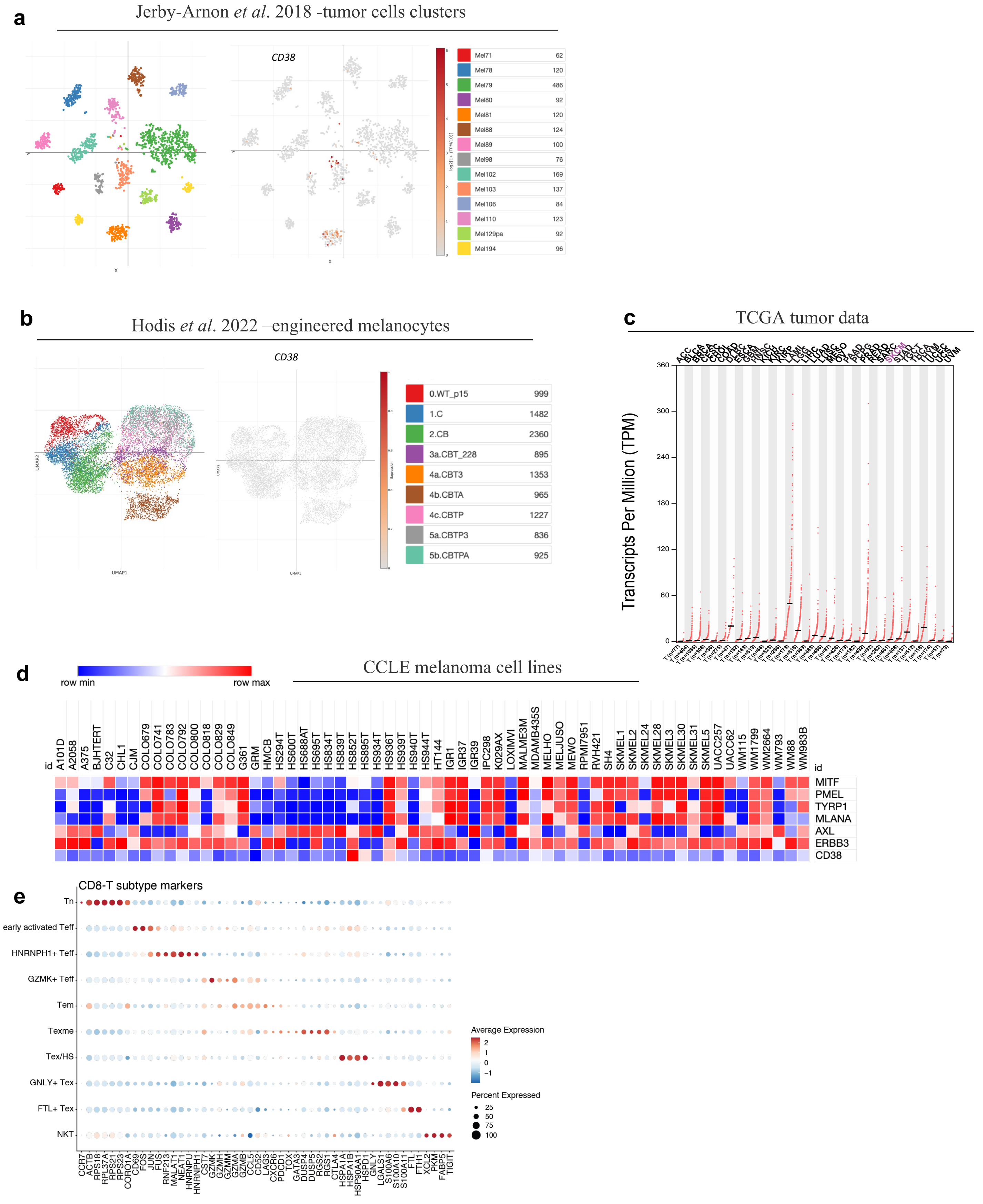


**Supplementary Data Fig. 2- a-b,** single cell RNAseq tSNE plots of melanoma tumour cells clusters and mapping of *CD38* expression in them [60,61]. **c,** CD38 expression in tumours from TCGA dataset versus normal tissue. Melanoma (SKCM) is highlighted in purple. **d,** *CD38* expression in melanoma cell lines from the CCLE database, clustered against proliferative melanoma (*MITF* high) versus neural crest like melanoma (*AXL* high). Supporting Extended Data Fig. 1k. **e,** T cell markers that were used for the classification of T cell clusters (Extended Data Fig. 2l).

Supplementary Data Figure 3

**a**

Gating strategy for Fig. 1i-j

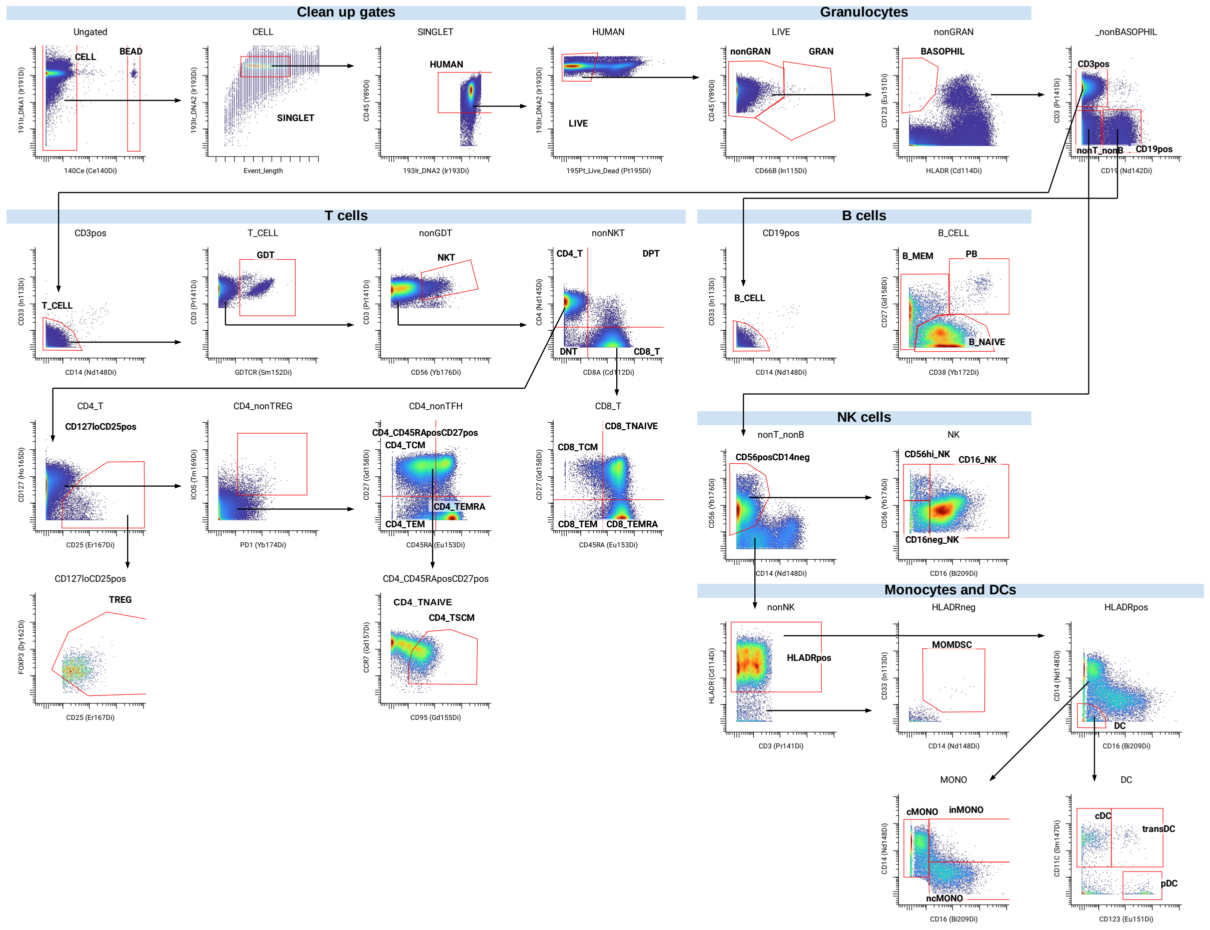


**Supplementary Data Fig. 3- a,** gating strategy for patient blood CD8+ T cells CyTOF analysis. Supporting Fig. 1i-j , Extended Data Fig. 2c-e.

### Supplementary Data Figure 4

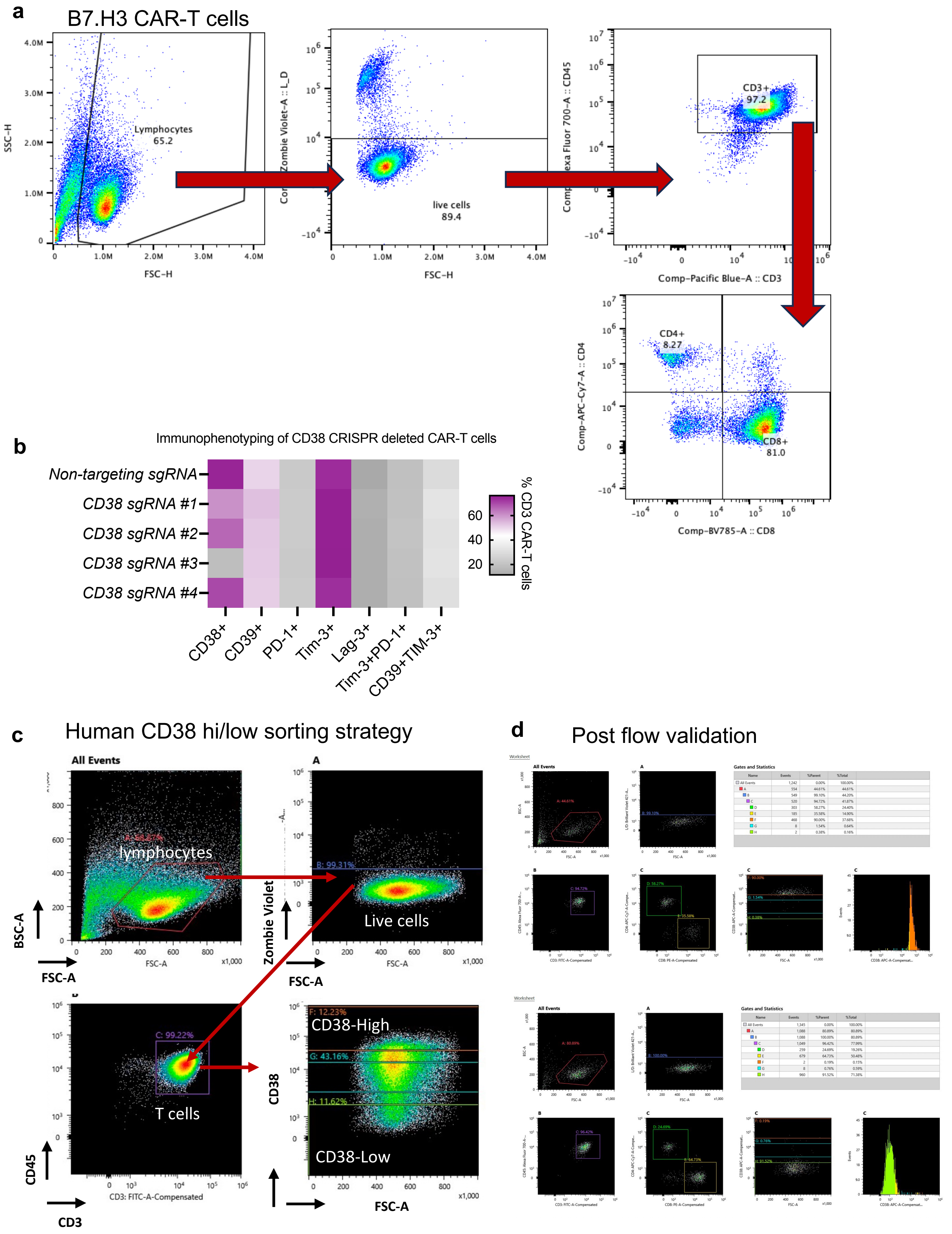


**Supplementary Data Fig. 4- a,** gating strategy for B7.H3 CAR-T cells. **b,** flow analysis of CD38 CRISPR edited B7.H3 CAR-T cells. **c-d,** B7.H3 CAR-T cells CD38 hi/low sorting strategy.

Supplementary Data Figure 5

PDOTS CD4+ and CD8 gating- CD8+CD38+/CD8+CD38+PD-1+/CD4+CD38+/CD4+CD38+PD-1+

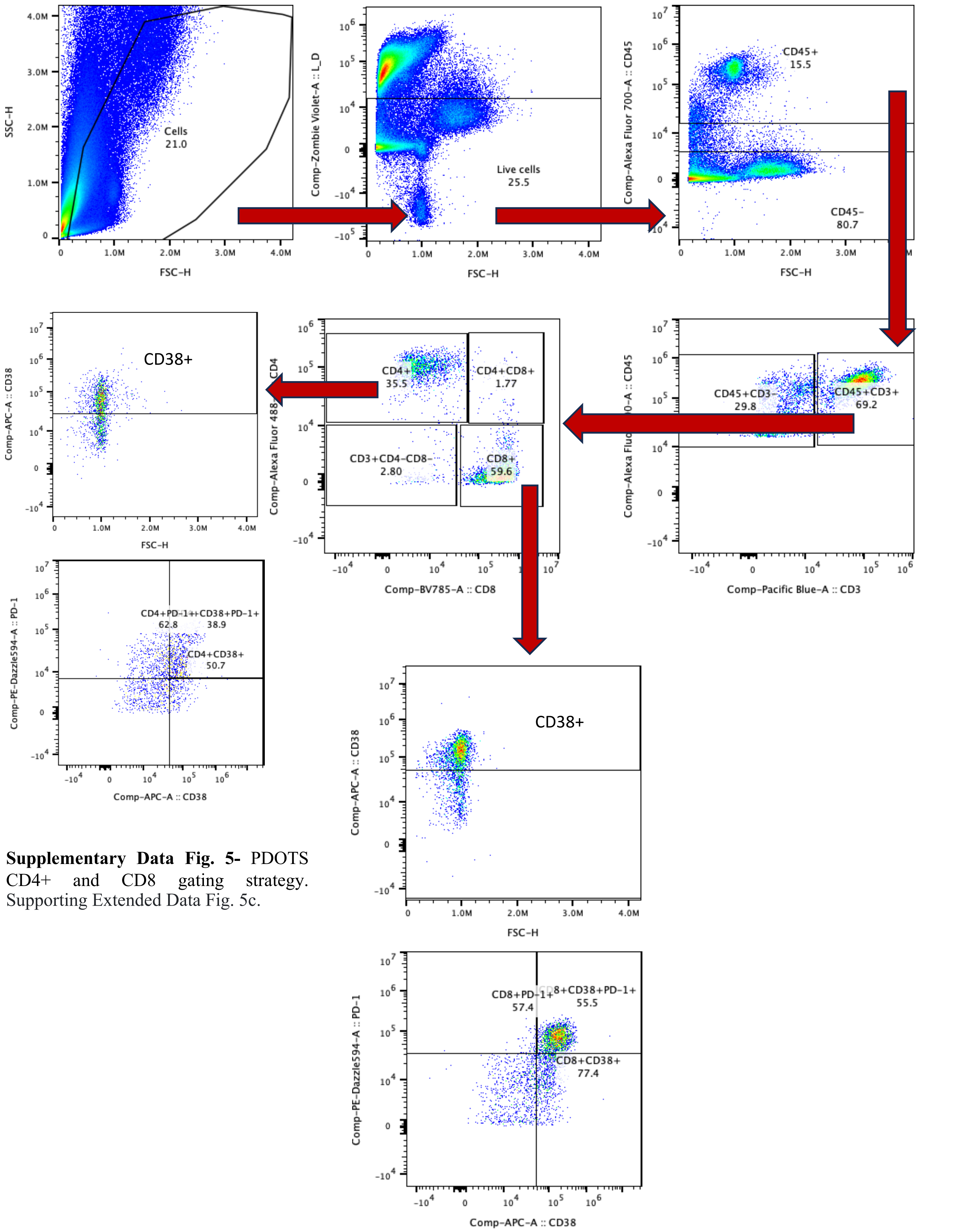


**Supplementary Data Fig. 5-** PDOTS CD4+ and CD8 gating strategy. Supporting Extended Data Fig. 5c.

Supplementary Data Figure 6

Mouse tumors gating- CD8+CD38+/ CD8+CD38+PD-1+/CD4+CD38+/ CD4+CD38+PD-1+

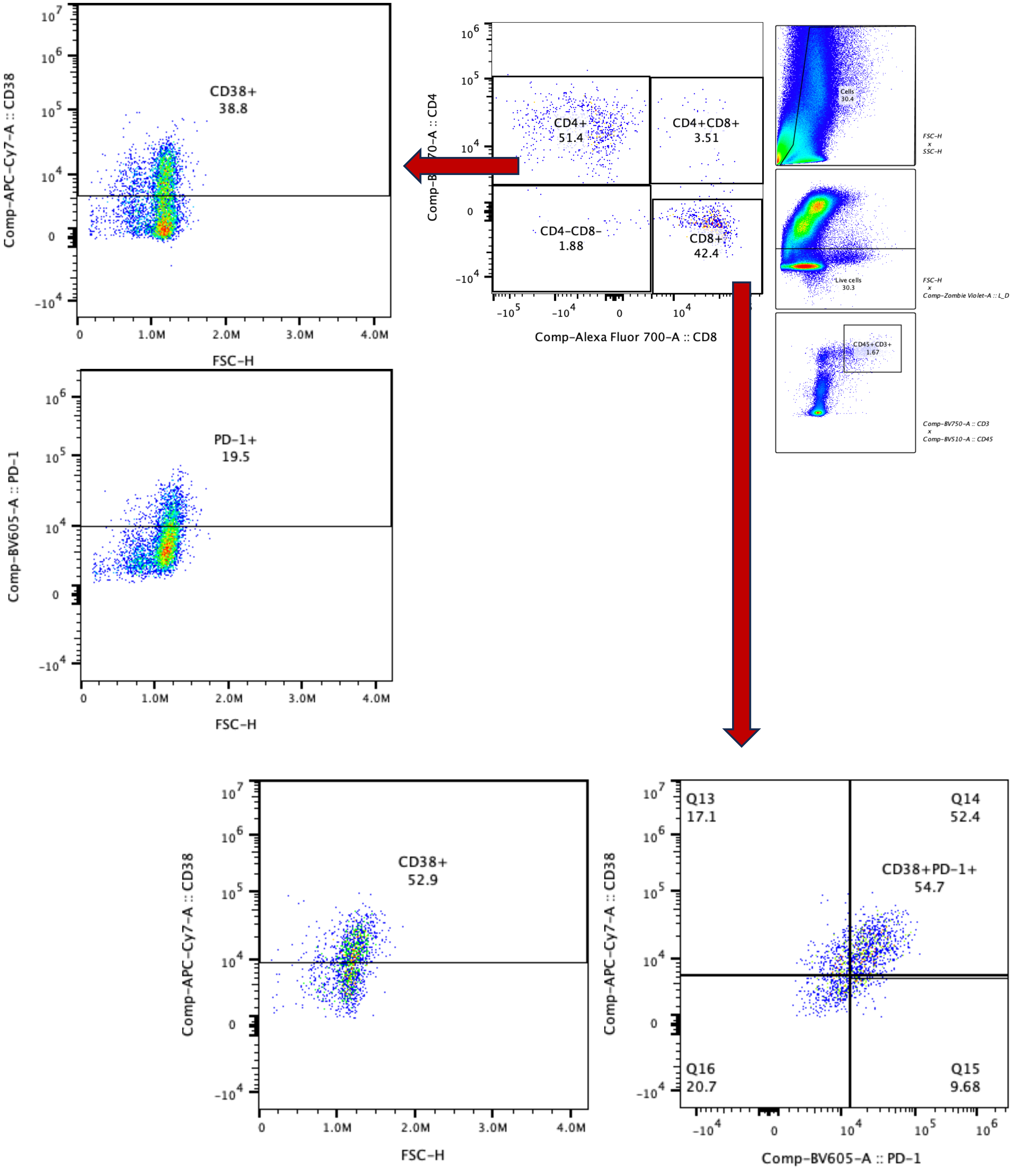


**Supplementary Data Fig. 6-** Mouse tumours gating strategy. Supporting Extended Data Fig. 6d.

Supplementary Data Figure 7

Mouse CD8+ isolated TILs

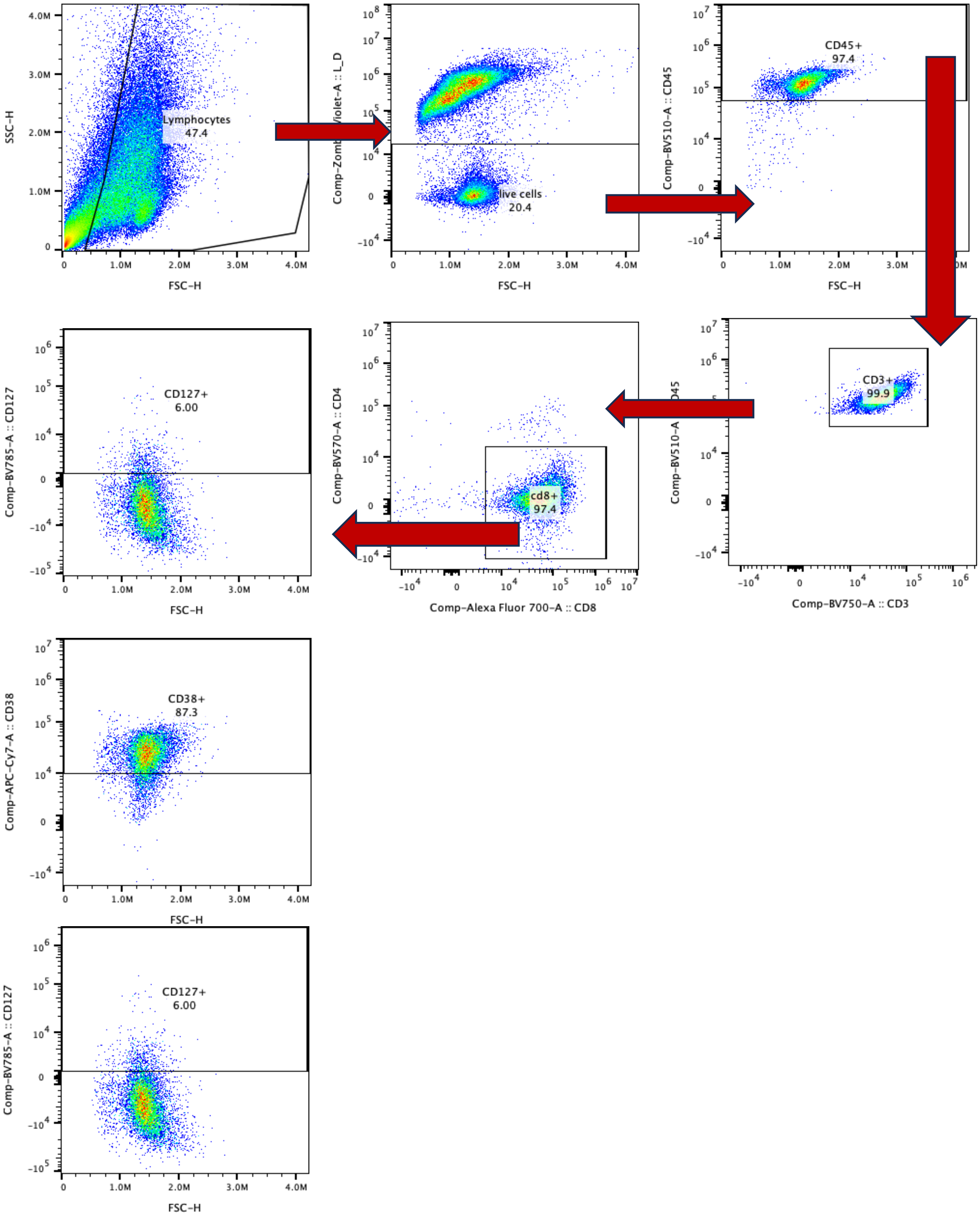


**Supplementary Data Fig. 7-** Mouse CD8+ isolated TILs gating strategy. Supporting Extended Data Fig. 6g-i

Supplementary Data Figure 8

**a** Mitochondrial stain gating

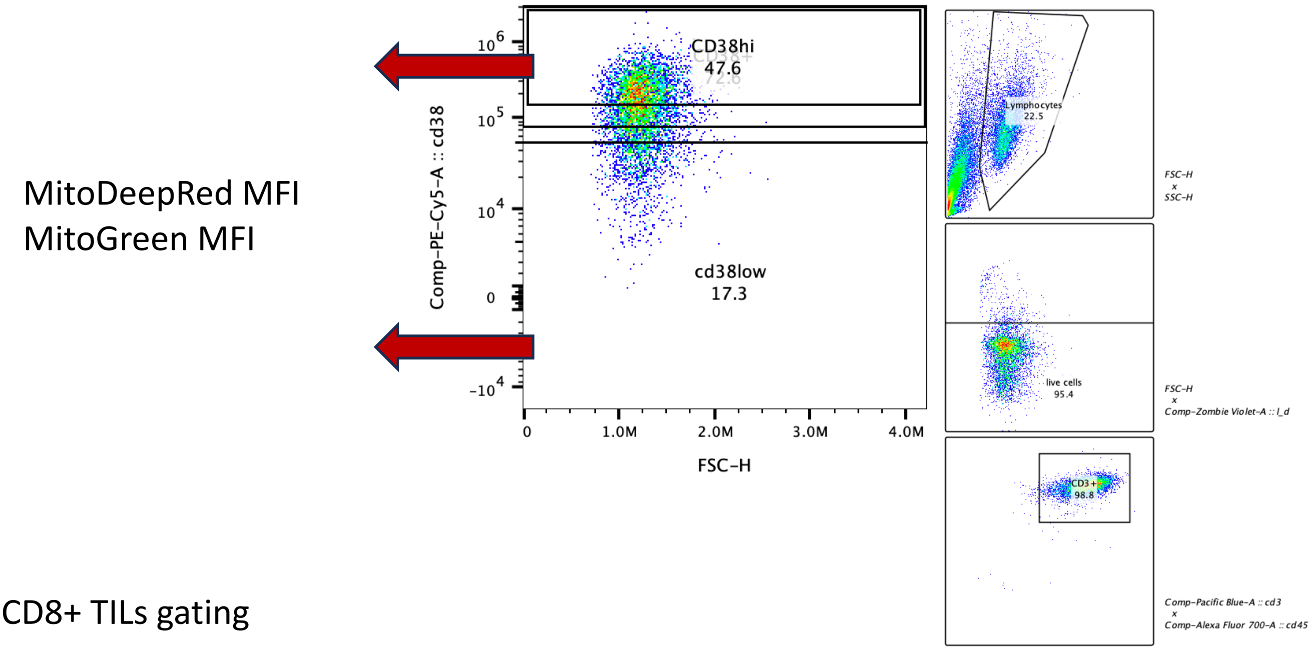


**b** CD8+ TILs gating

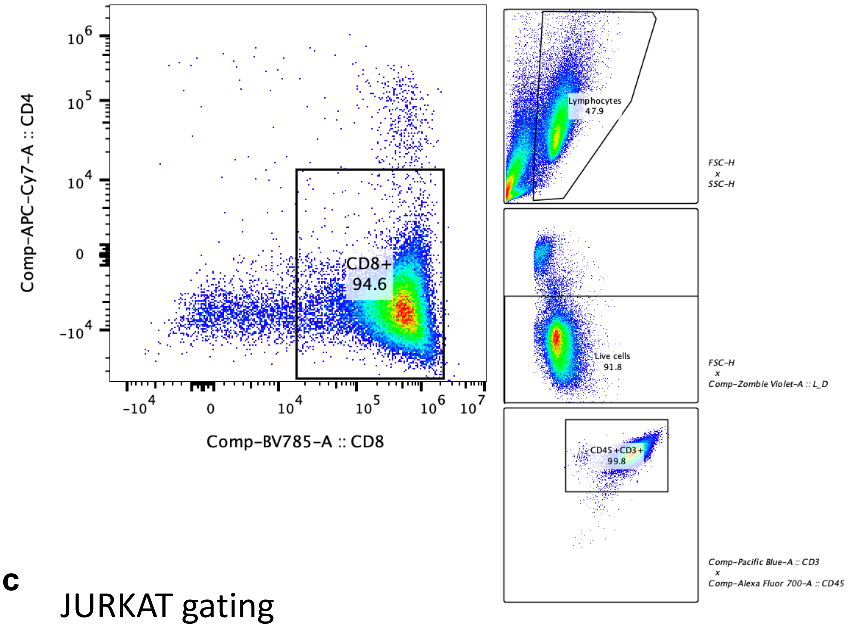


**c** JURKAT gating

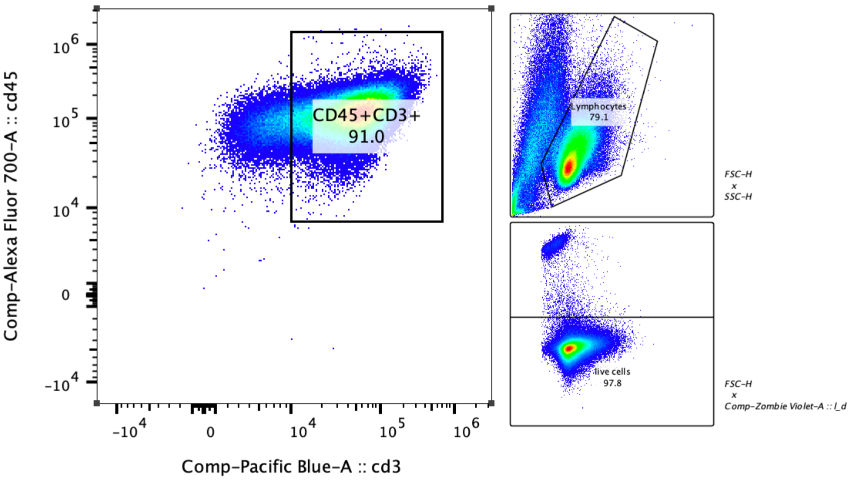


**Supplementary Data Fig. 8-** Mitochondrial stain gating strategy. Supporting Extended Data Fig. 7f-m.

### Supplementary Data Figure 9

a

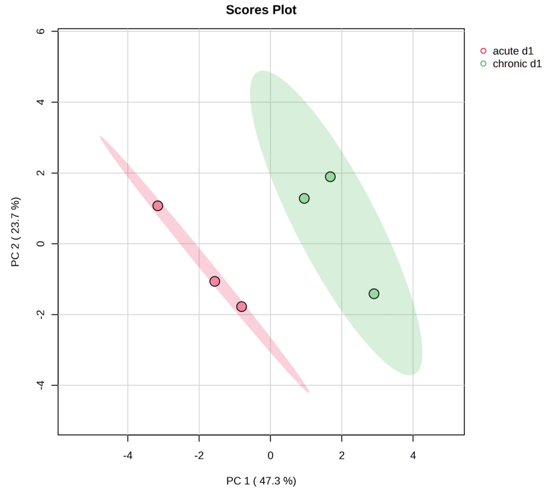


b

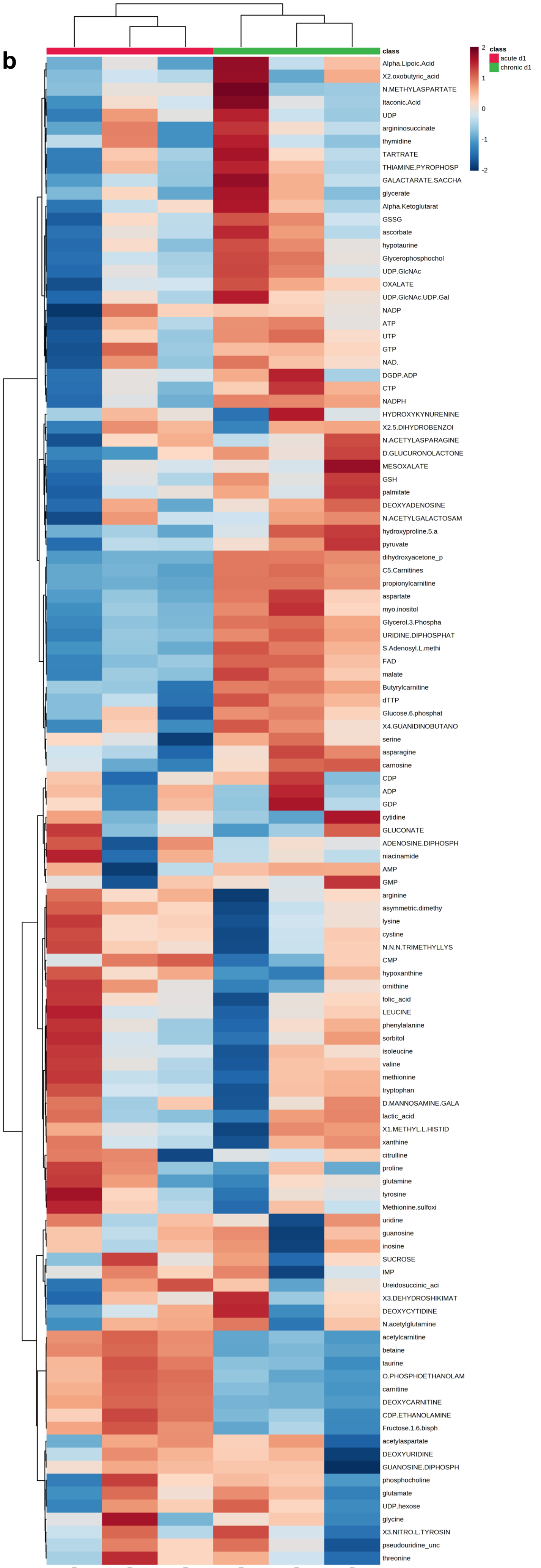


**Supplementary Data Fig. 9-** Intracellular metabolomics of acute versus chronically stimulated CAR-T cells. Supporting Extended Data Fig. 8c-d.

Supplementary Data Figure 10

**a**

#### Principle component analysis donor 1 and 2 intracellular metabolomics

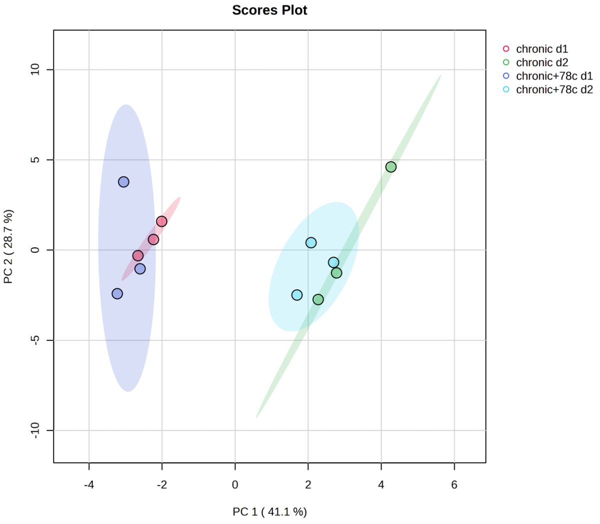


**b**

#### Principle component analysis donor 1 and 2 extracellular metabolomics

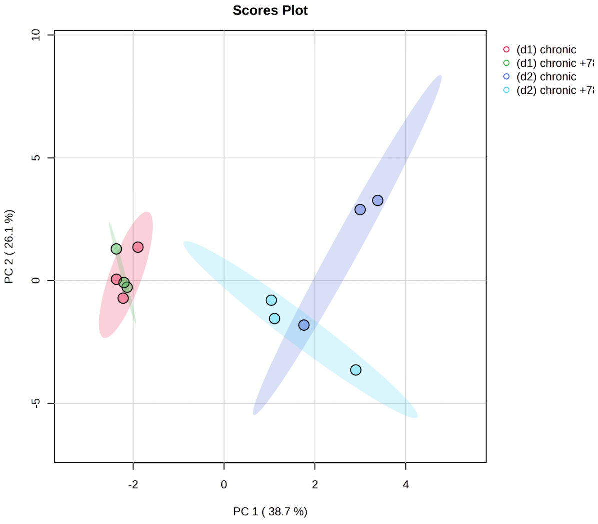


**Supplementary Data Fig. 10-12-** Intracellular and extracellular metabolomics of chronically stimulated CAR-T cells +/- CD38i. Supporting Fig. 4d-h, Extended Data Fig. 8e-i. 8l-o.

### Supplementary Data Figure 11

**a**

#### Intracellular metabolomics donor 1

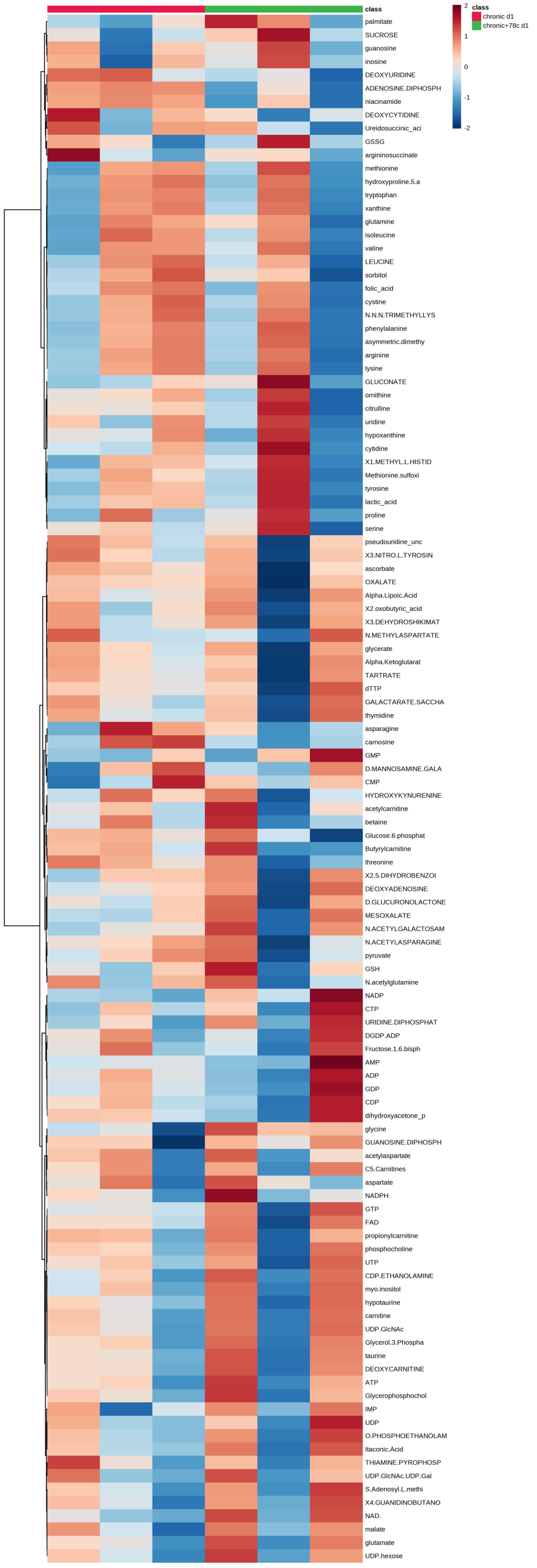


**b**

#### Intracellular metabolomics donor 2

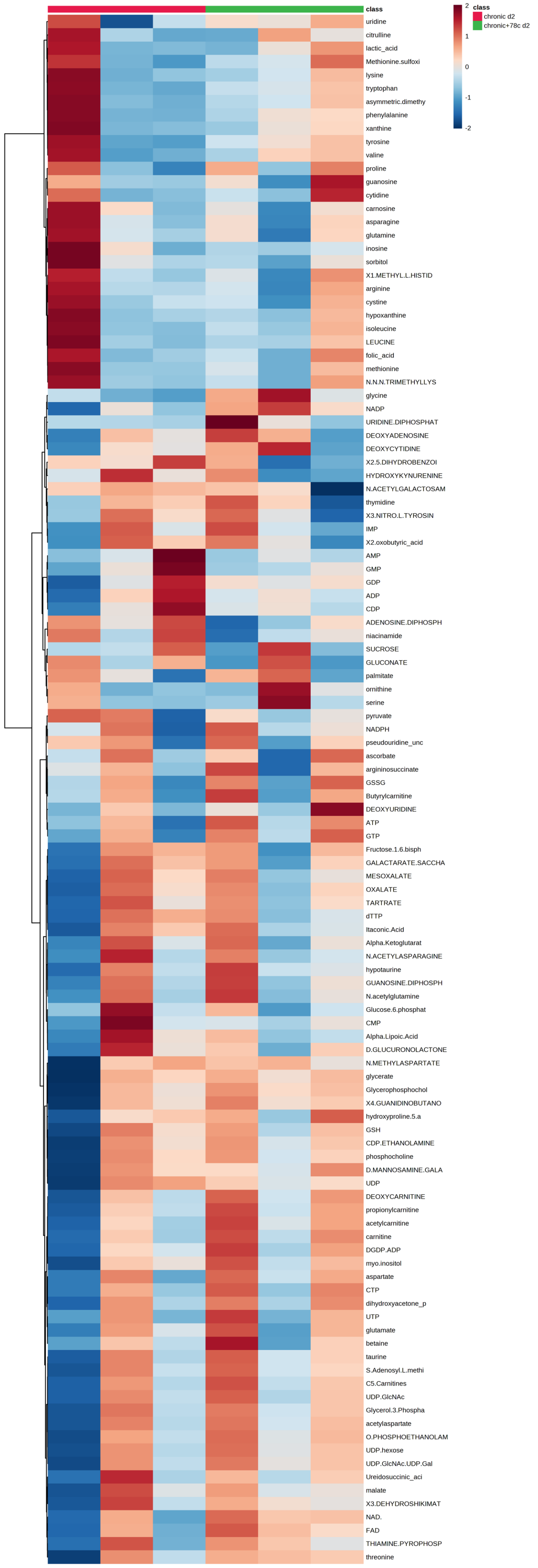

Supplementary Data Figure 12

**a**

#### Extracellular metabolomics donor 1

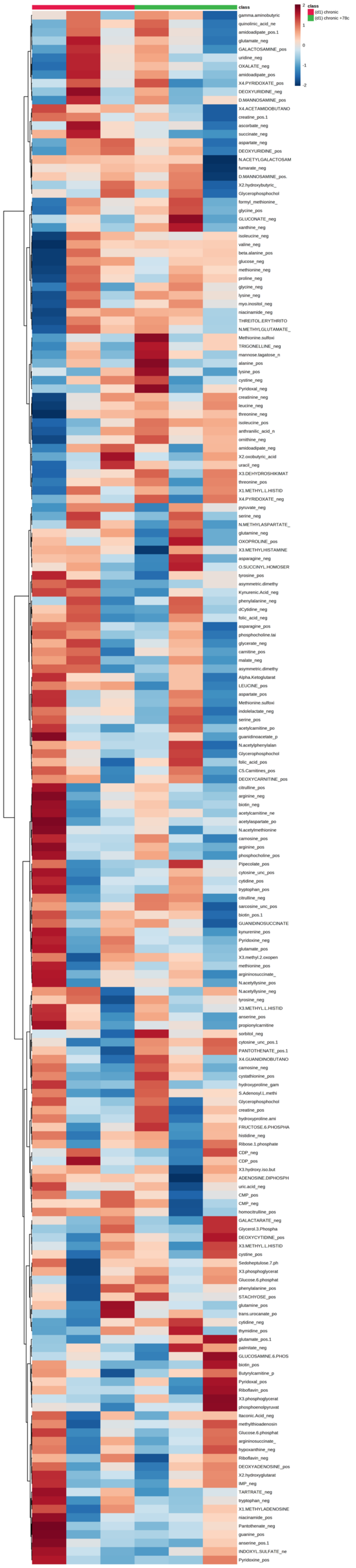

**b**

#### Extracellular metabolomics donor 2

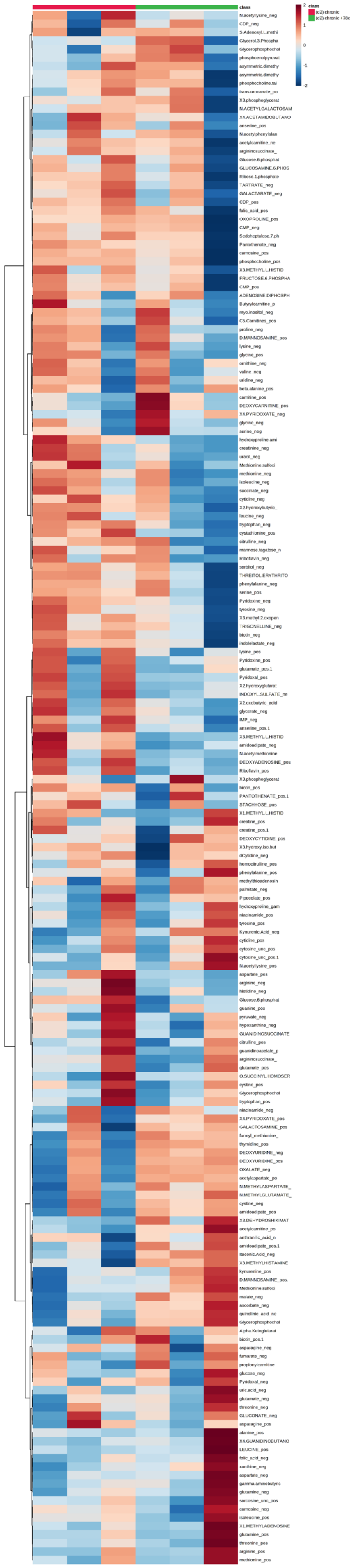
